# Microglia mediate synaptic plasticity induced by 10 Hz repetitive transcranial magnetic stimulation

**DOI:** 10.1101/2021.10.03.462905

**Authors:** Amelie Eichler, Dimitrios Kleidonas, Zsolt Turi, Maximilian Fliegauf, Matthias Kirsch, Dietmar Pfeifer, Takahiro Masuda, Marco Prinz, Maximilian Lenz, Andreas Vlachos

## Abstract

Microglia—the resident immune cells of the central nervous system—sense the activity of neurons and regulate physiological brain functions. They have been implicated in the pathology of brain diseases associated with alterations in neural excitability and plasticity. However, experimental and therapeutic approaches that modulate microglia function in a brain-region-specific manner have not been established. In this study, we tested for the effects of repetitive transcranial magnetic stimulation (rTMS), a clinically employed non-invasive brain stimulation technique, on microglia-mediated synaptic plasticity. 10 Hz electromagnetic stimulation triggered a release of plasticity-promoting cytokines from the microglia in organotypic brain tissue cultures, while no changes in microglial morphology or microglia dynamics were observed. Indeed, substitution of tumor necrosis factor alpha (TNFα) and interleukin 6 (IL6) preserved synaptic plasticity induced by 10 Hz stimulation in the absence of microglia. Consistent with these findings, *in vivo* depletion of microglia abolished rTMS-induced changes in neurotransmission in the medial prefrontal cortex (mPFC) of anesthetized mice. We conclude that rTMS affects neural excitability and plasticity by modulating the release of cytokines from microglia.

## Introduction

Despite its increasingly prevalent clinical use, e.g., for the treatment of patients with pharmaco-resistant depression, the cellular and molecular effects of repetitive transcranial magnetic stimulation (rTMS) remain poorly understood (Cirillo et al., 2017; Muller-Dahlhaus and Vlachos, 2013). It is widely known however, that the electric fields produced by non-invasive magnetic brain stimulation modulate neural excitability and synaptic transmission in cortical circuits (Pell et al., 2011). These changes require neural activity, i.e., action potential induction, and Ca^2+^-dependent signaling pathways, i.e., NMDA receptors and L-Type voltage-gated calcium channels (Lenz et al., 2015). Such changes are consistent with the long-term potentiation (LTP) of synaptic neurotransmission (Vlachos et al., 2012). It remains unclear however, how rTMS and the induction of “LTP-like” plasticity asserts positive effects in clinical settings.

Recent work suggests that microglia, the resident immune cells of the brain, sense and regulate synaptic transmission and plasticity (Badimon et al., 2020; Schafer et al., 2012). There is evidence that changes in network activity modulate microglia states (Pfeiffer et al., 2016; Stowell et al., 2019). These changes are reflected by the dynamic extension and retraction of microglial processes, by the formation of physical contacts between microglia and neurons, and by the activity-dependent secretion of plasticity-modulating factors (Akiyoshi et al., 2018; Cserep et al., 2020; Sheppard et al., 2019). Specifically, the plasticity-modulating effects of pro-inflammatory cytokines, such as tumor necrosis factor alpha (TNFα) and interleukin 6 (IL6), have been demonstrated in several experimental settings, both *in vitro* and *in vivo* (Heir and Stellwagen, 2020; Santello et al., 2011; Stellwagen and Malenka, 2006). These findings suggest that microglia play an important role in neural function and plasticity. Considering their role in pathological brain states (Graeber and Streit, 2010; Prinz et al., 2019; Prinz et al., 2021), it is interesting to theorize that the therapeutic effects of rTMS could—at least in part—depend on the modulation of microglia function. Indeed, evidence has been provided that rTMS affects microglial markers (Clarke et al., 2017; Cullen and Young, 2016; Li et al., 2021; Muri et al., 2020; Stevanovic et al., 2019). However, direct experimental evidence determining the role of microglia in rTMS-induced neural plasticity is currently not available.

To address the biological relevance of microglia in rTMS-induced synaptic plasticity, we depleted microglia from organotypic brain tissue cultures and the adult mouse brain and probed synaptic plasticity with a 10 Hz stimulation protocol. In turn, we tested for the effects of 10 Hz magnetic stimulation on the structural and functional properties of microglia and employed *in vitro* cytokine release assays to identify the plasticity promoting effects of microglial factors secreted after rTMS.

## Materials and Methods

### Ethics statement

Mice were maintained in a 12 h light/dark cycle with food and water available *ad libitum*. Every effort was made to minimize distress and pain of animals. All experimental procedures were performed according to German animal welfare legislation and approved by the competent authority (Regierungspräsidium Freiburg, G-20/154) appropriate animal welfare committee and the animal welfare officer of Albert-Ludwigs-University Freiburg, Faculty of Medicine (X-17/07K, X-17/09C, X-18/02C).

### Preparation of tissue cultures

Entorhino-hippocampal tissue cultures were prepared at postnatal day 3-5 from *C57BL/6J*, *HexB-tdTom* (Masuda et al., 2020), and *C57BL/6-Tg(TNFa-eGFP)* (Lenz et al., 2020) mice of either sex as previously described (Lenz et al., 2015). Cultivation medium contained 50% (v/v) MEM, 25% (v/v) basal medium eagle, 25% (v/v) heat-inactivated normal horse serum, 25 mM HEPES buffer solution, 0.15% (w/v) bicarbonate, 0.65% (w/v) glucose, 0.1 mg/ml streptomycin, 100 U/ml penicillin, and 2 mM glutamax. The pH was adjusted to 7.3. All tissue cultures were allowed to mature for at least 18 days in a humidified atmosphere with 5% CO2 at 35°C. Cultivation medium was replaced 3 times per week.

### Microglia depletion *in vitro* and *in vivo*

Tissue cultures were treated immediately after preparation (div 0) with the CSF-1R inhibitor PLX3397 (50 nM; #2501 Axon) for at least 18 days. Vehicle-only treated cultures (DMSO, 0.1 µl) served as age- and time-matched controls. For *in vivo* microglia depletion, we used the CSF-1R inhibitor BLZ945 (kindly provided by Novartis, Basel, Switzerland) dissolved in 20% (2-hydroxypropyl)-β-cyclodextrin (Sigma-Aldrich, Germany). A dose of 200 mg/kg bodyweight was applied by oral gavage in adult (8-weeks old) mice for 7 consecutive days as previously described (Hagemeyer et al., 2017; Masuda et al., 2020). No weight loss or any apparent signs of stress could be detected throughout the treatment period.

### Repetitive magnetic stimulation *in vitro*

Tissue cultures (≥ 18 days *in vitro*) were transferred to a 35 mm Petri Dish filled with pre-warmed standard extracellular solution containing (in mM): 129 NaCl, 4 KCl, 1 MgCl_2_, 2 CaCl_2_, 4.2 glucose, 10 HEPES, 0.1 mg/ml streptomycin, 100 U/ml penicillin, pH 7.4 adjusted with NaOH, osmolarity adjusted with sucrose to 380-390 mOsm. Cultures were stimulated using the Magstim Rapid^2^ stimulator (Magstim Company, UK) connected to a Double AirFilm^®^ Coil (coil parameters according to manufacturer’s description: average inductance = 12 μH; pulse rise time approximately 80 μs; pulse duration = 0.5 ms, biphasic; Magstim Company, UK) with a biphasic current waveform. Cultures were positioned approximately 1 cm under the center of the coil and oriented in a way that the induced electric field was parallel to the dendritic tree of CA1 pyramidal neurons. The stimulation protocol consisted of 900 pulses at 10 Hz (50% maximum stimulator output). Cultures were kept in the incubator for at least 2 h after stimulation before experimental assessment. Age- and time-matched control cultures were not stimulated, but otherwise treated identical to stimulated cultures (sham stimulation). For cytokine substitution experiments TNFα (5 ng/ml; #410-MT RD Systems) and IL6 (2.5 ng/ml; #406-ML RD Systems) were added to the stimulation medium.

### Repetitive magnetic stimulation *in vivo*

rTMS was carried out in adult (∼8 weeks old) urethane-anesthetized (1.25 g kg^−1^, intraperitoneal; 0.125 g kg^−1^, subcutaneous) *C57BL/6J* mice of either sex. The head was placed under the coil with the medial prefrontal cortex (mPFC) under the center. During the stimulation, the brain-to-coil distance was kept minimal, while brain-to-coil contact was avoided. Repetitive stimulation was performed at fixed intensities of 60% MSO (which corresponds to 90% motor threshold, see (Lenz et al., 2016)) using the same 10-Hz stimulation protocol described above. Control animals placed near the coil during stimulation were not stimulated but otherwise treated identically. All animals were transferred back to their cages with appropriate body temperature control and were held in anesthesia for 2 h under continuous surveillance. After the waiting period, ketamine/xylazin (100 mg kg^-1^ / 20 mg kg^-1^; i.p. application) was injected to achieve a suitable analgesia before rapid decapitation.

The brain was prepared as previously described (Lenz et al., 2021; Ting et al., 2018). After dissection, the brain was embedded in low-melting agar (1.8% w/v in PBS; Sigma Aldrich #A9517) and frontal sections (350 µm thickness) containing the mPFC were prepared using a Leica VT1200S vibratome with a cutting angle of 15°. The brain was cut in NMDG-aCSF [containing (in mM) 92 NMDG, 2.5 KCl, 1.25 NaH_2_PO_4_, 30 NaHCO_3_, 20 HEPES, 25 glucose, 2 thiourea, 5 Na-ascorbate, 3 Na-pyruvate, 0.5 CaCl_2_, and 10 MgSO_4_, (pH = 7.3–7.4)] at approximately 0°C. After cutting slices were recovered in NMDG-aCSF at 34°C. Sodium spike-in was performed according to a previously established protocol that is suitable for ∼8 week old animals (Lenz et al., 2021; Ting et al., 2018). After recovery, we transferred the slices to a holding chamber [holding-aCSF; containing (in mM) 92 NaCl, 2.5 KCl, 1.25 NaH_2_PO_4_, 30 NaHCO_3_, 20 HEPES, 25 glucose, 2 thiourea, 5 Na-ascorbate, 3 Na-pyruvate, 2 CaCl_2_, and 2 MgSO_4_, (pH = 7.3–7.4)] at room temperature in which slices were maintained at least half an hour until electrophysiological assessment.

### Propidium iodide staining

Tissue cultures were incubated with propidium iodide (PI, 5 μg/ml; #P3566 Invitrogen) for 2 h, washed in phosphate buffered saline (PBS) and fixed as described below. Cultures treated for 4 h with NMDA (50 μg/ml; #0114 Tocris) served as positive controls in these experiments. Cell nuclei were stained with DAPI, sections were mounted on microscope slides and confocal images were acquired as described below.

### Immunostaining and imaging

Tissue cultures were fixed overnight in a solution of 4% (w/v) paraformaldehyde (PFA) in PBS (prepared from 16% PFA stocks in phosphate buffered saline according to manufacturer’s instruction; #28908 Thermo Scientific). After fixation, cultures were washed in PBS (0.1 M, pH 7.4) and consecutively incubated for 1 h with 10% (v/v) normal goat serum (NGS) in 0.5% (v/v) Triton X-100 containing PBS to reduce nonspecific staining and to increase antibody penetration. Subsequently, cultures were incubated overnight at 4°C with rabbit anti-Iba1 (1:1000; #019-19741 Fujifilm Wako) in PBS with 10% NGS and 0.1% Triton X-100. Sections were washed and incubated overnight at 4°C with appropriate Alexa Fluor^®^ dye-conjugated secondary antibodies (1:1000, donkey anti-rabbit Alexa Fluor 488 or 647; #A-21206 or #A-32795 Invitrogen) in PBS with 10% NGS or NHS, 0.1% Triton X-100. For post-hoc visualization of patched pyramidal cells, Streptavidin Alexa Fluor 488 or 633 (Streptavidin A488, 1:1000; #S32354 Invitrogen; Streptavidin A633, 1:1000; #S21375 Invitrogen) was added to the secondary antibody incubation solution. DAPI nuclear stain (1:2000 in PBS for 20 minutes; #62248 Thermo Scientific) was used to visualize cytoarchitecture. Cultures were washed, transferred onto glass slides and mounted for visualization with DAKO anti-fading mounting medium (#S302380-2 Agilent).

A Leica SP8 laser-scanning microscope equipped with a 20x multi-immersion (NA 0.75; Leica), a 40x oil-immersion (NA 1.30; Leica) and a 63x oil-immersion objective (NA 1.40; Leica) was used for confocal image acquisition. Images for analysis of microglia cell density (Figure 1) and images of propidium iodide stainings (Figure 3) were acquired with a 20x objective at 0.75x optical zoom (resolution: 512 x 512 px). Image stacks for spine density and spine volume analysis (Figure 6) were acquired with a 63x oil-immersion objective at 5.0x optical zoom (resolution: 1024 x 1024, Δz = 0.22 µm at ideal Nyquist rate). Image stacks of Iba1 stained *HexB-tdTom* cultures (Figure S1) were acquired with a 20x objective at 2.0x optical zoom (resolution: 512 x 512 px). Image stacks of Iba1 stained acute cortical slices (Figure 11) were acquired using a 40x oil-immersion objective at 2.0x optical zoom (resolution 1024×1024, Δz = 1 µm). Laser intensity and detector gain were set to achieve comparable overall fluorescence intensity throughout stacks between all groups in each experimental setting.

**Figure 1:**
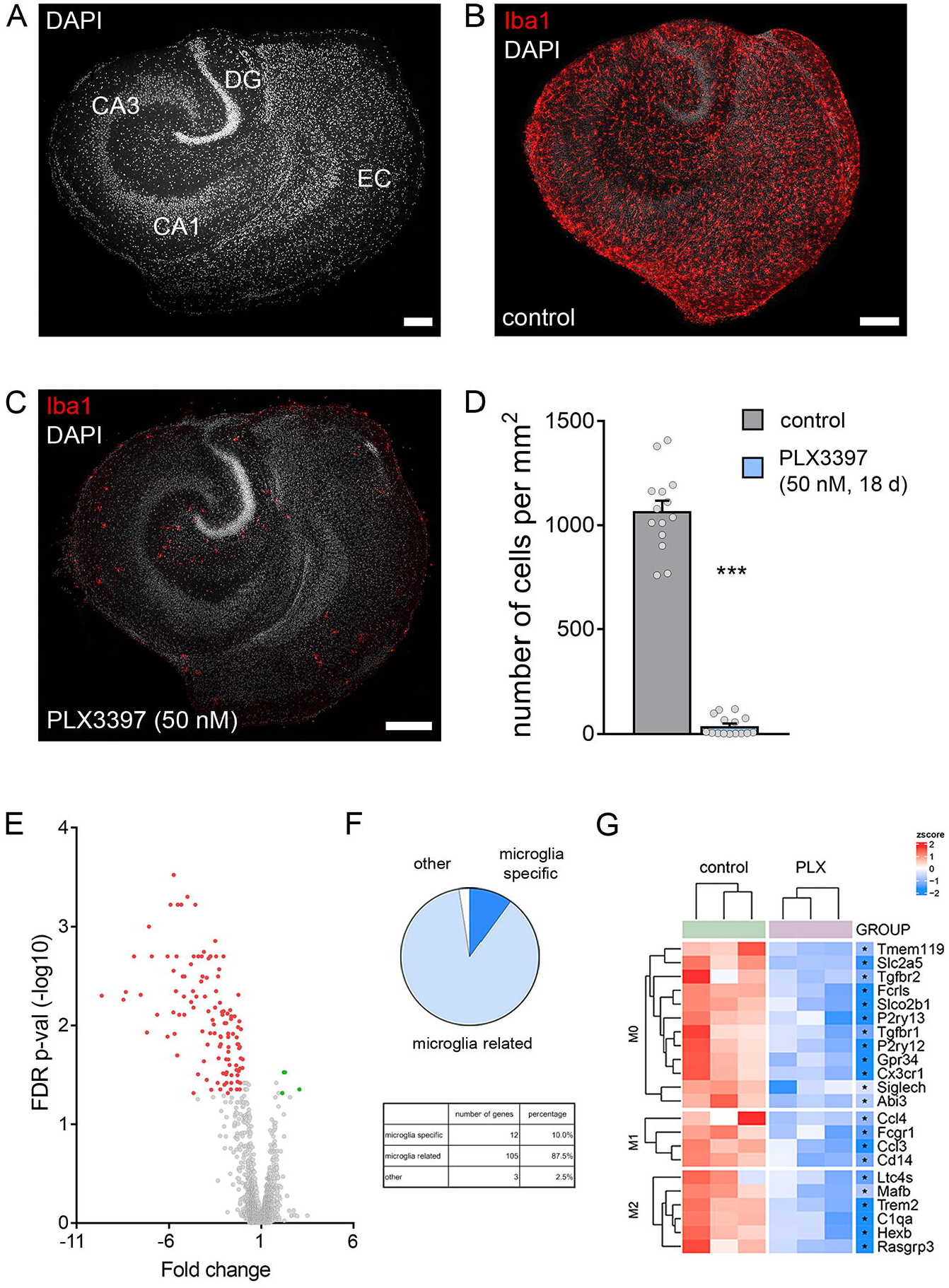
PLX3397 depletes microglia in organotypic tissue cultures. (A) Entorhino-hippocampal tissue culture stained with DAPI nuclear stain (EC, entorhinal cortex; DG, dentate gyrus). Scale bar, 200 µm. (B, C) Representative examples of tissue cultures stained for the microglial marker Iba1. Note homogenous distribution of microglia in the control culture and almost complete depletion of microglia following PLX3397 treatment (50 nM, 18 days). Scale bars, 200 µm. (D) Microglia cell counts in the respective groups (n_control_ = 14 cultures, n_PLX(50nM)_ = 15 cultures; Mann-Whitney test, U = 0). (E-G) Affymetrix^®^ Microarray analysis of control cultures and cultures treated with PLX3397. (E) Volcano plot shows fold changes and FDR p-values of analyzed transcripts. Significantly upregulated transcripts are indicated in green, significantly downregulated transcripts are indicated in red. (F) Classification of differentially expressed transcripts. 97.5% of the differentially expressed transcripts are microglia-specific or microglia-related. (G) Hierarchical clustering of differentially expressed gene sets characteristic of M0-, M1-, and M2-classified microglia. Each sample consisted of 3 pooled cultures (n = 3 samples in each group). Individual data points are indicated by colored dots. Values represent mean ± s.e.m (***p < 0.001).

**Figure 2:**
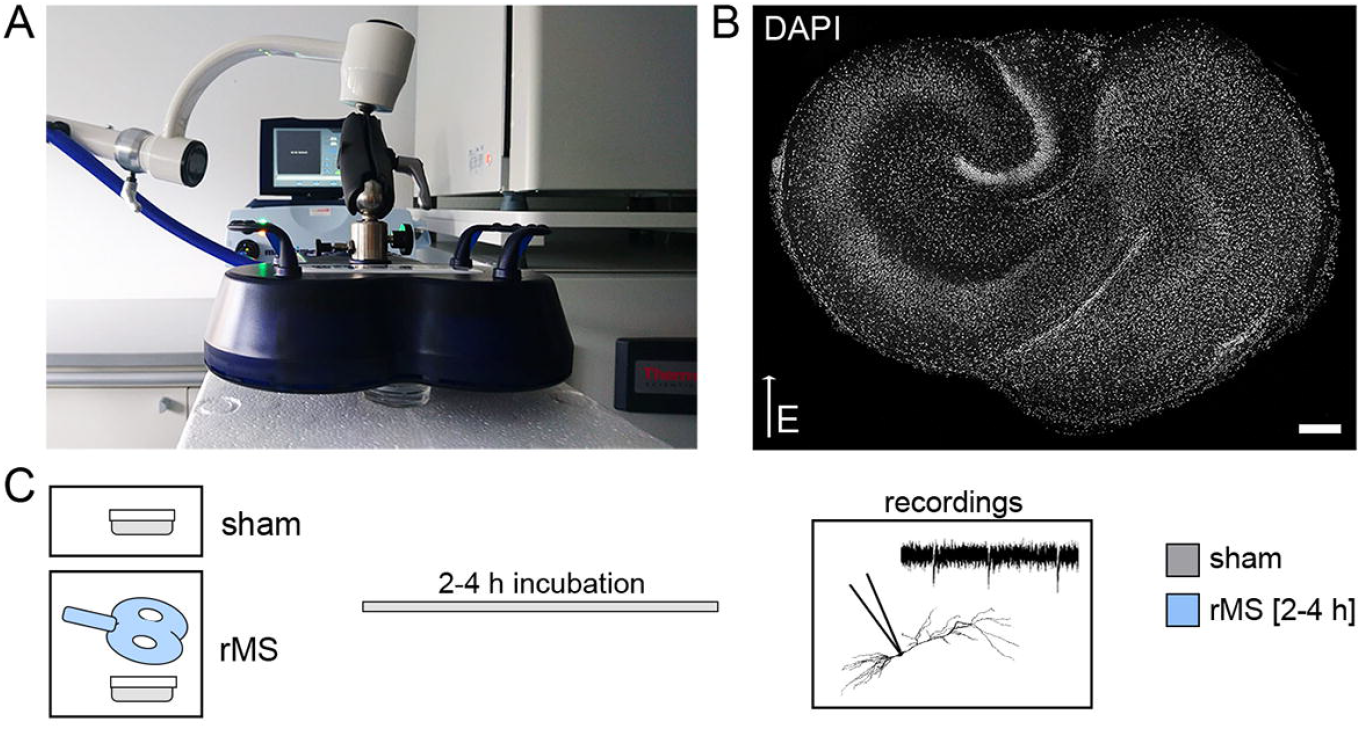
10 Hz repetitive magnetic stimulation in organotypic tissue cultures. (A-B) Entorhino-hippocampal tissue cultures (DAPI nuclear stain, B) were stimulated with a 70 mm outer wing diameter figure of eight coil (Magstim Company, UK). Filter inserts carrying 2–6 tissue cultures were placed in a Petri Dish below the coil. The orientation within the electromagnetic field and distance to the coil was kept constant in all experiments (DAPI nuclear stain). Scale bar, 200 µm. (C) Sham stimulated cultures were not stimulated but otherwise treated equally. After stimulation, cultures were kept in the incubator for 2-4 h before further experimental procedures like patch-clamp recordings.

**Figure 3:**
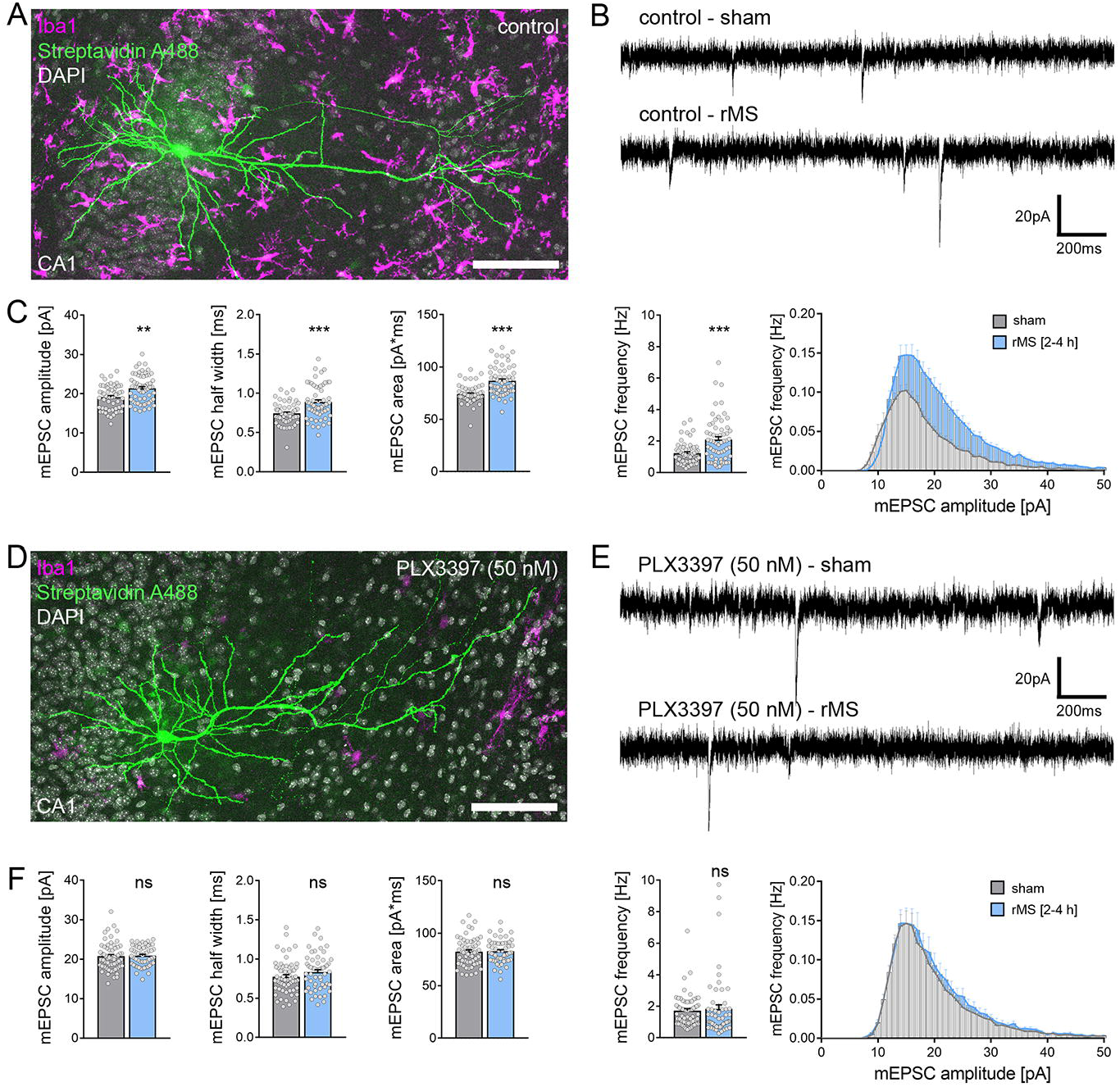
CA1 pyramidal neurons in microglia-depleted tissue cultures do not express excitatory synaptic plasticity induced by 10 Hz repetitive magnetic stimulation (rMS) (A) Example of a recorded and posthoc stained CA1 pyramidal neuron in a tissue culture stained with the microglial marker Iba1. Scale bar, 100 µm. (B, C) Sample traces and group data of AMPA receptor-mediated miniature excitatory postsynaptic currents (mEPSCs) recorded from CA1 pyramidal neurons in sham-stimulated and 10 Hz rMS-stimulated cultures 2-4 h after stimulation (n_control-sham_ = 56 cells, n_control-rMS_ = 60 cells; Mann-Whitney test, U_amplitude_ = 1091, U_half width_ = 1042, U_area_ = 956, U_frequency_ = 651; RM two-way ANOVA followed by Sidak’s multiple comparisons test for amplitude-frequency-plot). (D) Example of a recorded and posthoc stained CA1 pyramidal neuron in a PLX3397 treated, microglia-depleted tissue culture stained with the microglial marker Iba1. Scale bar, 100 µm. (E, F) Representative traces and group data of AMPA receptor-mediated mEPSCs recorded from CA1 pyramidal cells in sham-stimulated and 10 Hz rMS-stimulated microglia-depleted tissue cultures 2-4 h after stimulation (n_PLX3397-sham_ = 61 cells, n_PLX3397-rMS_ = 55 cells; Mann-Whitney test; RM two-way ANOVA followed by Sidak’s multiple comparisons test for amplitude-frequency-plot). Individual data points are indicated by grey dots. Values represent mean ± s.e.m (** p < 0.01, *** p < 0.001; ns, not significant differences).

### Live-cell imaging

Live-cell imaging of tissue cultures was performed at a Zeiss LSM800 microscope equipped with a 10x water-immersion (NA 0.3; Carl Zeiss) and a 40x water-immersion objective (NA 1.0; Carl Zeiss). Filter membranes with 2 to 6 cultures were placed in a 35 mm Petri Dish containing pre-oxygenated imaging solution consisting of 50% (v/v) MEM, 25% (v/v) basal medium eagle, 50 mM HEPES buffer solution (25% v/v), 0.65% (w/v) glucose, 0.15% (w/v) bicarbonate, 0.1 mg/ml streptomycin, 100 U/ml penicillin, 2 mM glutamax and 0.1 mM trolox. The cultures were kept at 35°C during the imaging procedure.

Live-cell imaging of homozygous (*HexB^tdT/tdT^*) and heterozygous (*HexB^tdT/+^*) cultures prepared from *HexB-tdTom* transgenic animals was performed to assess microglia morphology after rMS. Cultures were stimulated as described above (rMS and sham stimulation) and imaging was started immediately in imaging solution under continuous oxygenation (5% CO_2_ / 95% O_2_). For 3 hours, every 2 minutes a z-stack of the same cell was recorded using a 40x water-immersion objective with Δz = 1 μm at ideal Nyquist rate and an optical zoom of 1.0x (resolution 512 x 512 px, 2x line average). Laser intensity and detector gain were initially set and were kept constant over image acquisition time.

Live-cell imaging of *C57BL/6-Tg(TNFa-eGFP)* cultures was performed to monitor TNFα expression after rMS as an indicator of neuroinflammation. Cultures were stimulated as described above (rMS and sham stimulation) and kept in the incubator after stimulation. After 3 hours a z-stack of each culture was recorded using a 10x water-immersion objective with Δz = 6.3 μm at ideal Nyquist rate and an optical zoom of 0.5x (resolution 1024 x 1024 px). Laser intensity and detector gain were initially set to keep the fluorescent signal in a dynamic range throughout the experiment and were kept constant.

Confocal image stacks were stored as .czi files.

### Transcriptome Microarray

Tissue cultures that were cultivated on one filter membrane (3 cultures) were transferred as one sample into RLT buffer (QIAGEN) and RNA was isolated according to the manufacturer’s instructions (RNeasy Plus Micro Kit; #74034 QIAGEN). RNA was eluted in 50 µl water and precipitated in 0.75 M ammonium-acetate and 10 µg glycogen (#R0551 Thermo Scientific) by adding 125 µl ethanol (100%). Samples were incubated at −80°C overnight and consecutively centrifuged for 30 minutes at 4°C. Pellets were washed with 70% ethanol, centrifuged again and dried. Finally, pellets were dissolved in water for further processing. RNA concentration and integrity was consecutively analyzed by capillary electrophoresis using a Fragment Analyser (Advanced Analytical Technologies, Inc., USA). RNA samples with RNA quality numbers (RQN) > 8.0 were further processed with the Affymetrix WT Plus kit and hybridized to Clariom S mouse arrays as described by the manufacturer (Thermo Fisher, Germany). Briefly, labeled fragments were hybridized to arrays for 16 h at 45°C, 60 rpm in a GeneChip™ Hybridization Oven (Thermo Fisher, Germany). After washing and staining, the arrays were scanned with the Affymetrix GeneChip Scanner 3000 7G (Thermo Fisher, Germany). CEL files were produced from the raw data with Affymetrix GeneChip Command Console Software Version 4.1.2 (Thermo Fisher, Germany).

### Cytokine detection assay

To analyze protein release upon rMS, cultures were stimulated on incubation medium in interface configuration with three cultures grown on one filter membrane. Three hours after stimulation, both incubation medium (for detection of protein release) and tissue cultures (for gene expression analysis) were collected and frozen in liquid nitrogen until further processing.

For cytokine detection, a V-Plex Proinflammatory Panel 1 (mouse) Kit Plus (#K15048G Mesoscale Discovery) was used. The collected incubation medium was diluted 1:1 in diluent provided with the kit. Protein detection was performed according to the manufacturer’s instructions. A pre-coated plate with capture antibodies on defined spots was incubated with the diluted samples overnight. After washing, samples were incubated for over night with a solution containing electrochemiluminescent MSD SULFO-TAG detection antibodies (Mesoscale Discovery; Antibodies: Anti-ms TNFα Antibody #D22QW, Anti-ms IL6 Antibody #D22QX, Anti-ms CXCL1 Antibody #D22QT). After washing, samples were measured with a MESO QuickPlex SQ 120 instrument (Mesoscale Discovery). The respective protein concentrations were determined using the MSD DISCOVERY WORKBENCH software (Mesoscale Discovery).

### RNA Isolation and Quantitative reverse transcription PCR (RT-qPCR)

RNA isolation for qPCR analysis was performed as follows. Tissue cultures that were cultivated on one filter membrane (3 cultures) were transferred as one sample into RNA Protection buffer (New England Biolabs) and RNA was isolated according to the manufacturer’s instructions (Monarch^®^ Total RNA Miniprep Kit; #T2010S New England Biolabs). As a quality control, the RIN (RNA integrity number) value of RNA isolated from tissue culture was determined using the Agilent RNA 6000 Pico Kit (#5067-1513 Agilent) with a 2100 Bioanalyzer (#G2939BA Agilent; Mean RIN value: 8.9). Purified RNA was consecutively reverse transcribed (RevertAid RT Kit; #K1691 Thermo Scientific). cDNA was diluted in water to a final concentration of 3 ng/ml. RT-qPCR was performed using a C1000 Touch Thermal Cycler (BIO-RAD) and the CFX 384 Real-Time PCR system (BIO-RAD). 13.5 ng target cDNA diluted in TaqMan Gene Expression Master Mix (#4369016 Applied Biosystems) were amplified using standard TaqMan gene expression assays (Applied Biosystems; Assay-IDs: *Gapdh*: Mm99999915_g1; *Tnf*: Mm00443258_m1; *Il6*: Mm00446190_m1; *Cxcl1*: Mm04207460_m1). The RT-qPCR protocol was performed as follows: 1 cycle of 50°C for 2 min, 1 cycle of 95°C for 10 min, 40 cycles of 95°C for 15 s and 60°C for 1 min. Of each sample three technical replicates were used and no amplification was detected in non-template controls. Amplification curves were excluded from further analysis if efficiency values were less than 90 or exceeded 110 according to automated calculation by the Bio-Rad CFX Maestro software package. Data were exported and stored on a computer as .pcrd-files.

### Whole-cell patch-clamp recordings

Whole-cell patch-clamp recordings were carried out at 35°C (3-6 neurons per culture). Patch pipettes contained (in mM) 126 K-gluconate, 10 HEPES, 4 KCl, 4 ATP-Mg, 0.3 GTP-Na_2_, 10 PO-Creatine, 0.3% (w/v) biocytin (pH 7.25 with KOH, 290 mOsm with sucrose), having a tip resistance of 4-6 MΩ. Pyramidal neurons were visually identified using a LN-Scope (Luigs & Neumann) equipped with an infrared dot-contrast and a 40x water-immersion objective (NA 0.8; Olympus). Electrophysiological signals were amplified using a Multiclamp 700B amplifier, digitized with a Digidata 1550B digitizer, and visualized with the pClamp 11 software package.

For whole-cell patch-clamp recordings of CA1 pyramidal neurons in tissue cultures, the bath solution contained (in mM) 126 NaCl, 2.5 KCl, 26 NaHCO_3_, 1.25 NaH_2_PO_4_, 2 CaCl_2_, 2 MgCl2, and 10 glucose (aCSF) and was oxygenated continuously (5% CO_2_ / 95% O_2_). Spontaneous excitatory postsynaptic currents (sEPSCs) of CA1 pyramidal neurons were recorded in voltage-clamp mode at a holding potential of −60 mV. Series resistance was monitored before and after each recording and recordings were discarded if the series resistance reached ≥ 30 MΩ. For mEPSC recordings of CA1 pyramidal neurons, D-APV (10 μM; #ab120003 Abcam), (-)-Bicuculline-Methiodide (10 µM; #ab120108 Abcam) and TTX (0.5 μM; #18660-81-6 Biotrend) were added to the external solution.

Whole-cell patch-clamp recordings of superficial (layer 2/3) cortical pyramidal neurons in acute mouse brain slices were carried out in a bath solution containing holding-aCSF. For sEPSC recordings, layer 2/3 pyramidal neurons were held at −70 mV in voltage-clamp mode.

For recording of intrinsic cellular properties in current-clamp mode, pipette capacitance of 2.0 pF was corrected and series resistance was compensated using the automated bridge balance tool of the MultiClamp commander. I-V-curves were generated by injecting 1 s square pulse currents starting at −100 pA and increasing in 10 pA steps (sweep duration: 2 s). Series resistance was monitored, and recordings were discarded if the series resistance reached ≥ 30 MΩ.

### Multi-scale modeling

A 3-dimensional mesh model was created with two compartments, i.e., bath solution and organotypic tissue culture, using the finite element method and the program Gmsh (4.8.4). Local mesh resolution was increased from 0.01 to 0.004 units in the CA1 region of the culture, i.e., region of interest (ROI), where neurons were placed. The final mesh consisted of 3.55 × 10^6^ nodes and 2.11 × 10^7^ tetrahedrons. The mean tetrahedron edge length was 5.6 µm in the ROI. The physical dimensions of the mesh model were adapted from the *in vitro* setting.

The coil-to-culture distance was kept at 10 mm and the coil was positioned above the culture. Electrical conductivities for the bath and culture were 1.654 S m^-1^ and 0.275 S m^-1^, respectively. The rate of change of the coil current was set to 1.4 A µs^-1^ at 1% MSO and it was scaled up to 50% MSO. Macroscopic electric field simulations were performed using SimNIBS (3.2.4) and Matlab (2020b). A validated 70 mm MagStim figure-of-eight coil was used in all simulations.

For multi-scale modeling, we used the Neuron Modeling for TMS (NeMo-TMS) framework to study the biological responses of CA1 pyramidal neurons to biphasic single pulse TMS and rTMS (Shirinpour et al., 2021). Axonal morphology was adopted from an example cell (Shirinpour et al., 2021). For all neurons, we implemented the generalized version of the Jarsky model (Shirinpour et al., 2021).

We extracted the membrane potentials and voltage-gated calcium ‘influx’ from the somatic and dendritic compartments (Shirinpour et al., 2021). We analyzed the number of action potentials, calcium spikes and their peak values. Simulations were run on a high performance computer in the state of Baden-Württemberg, Germany (bwHPC).

### Quantification and statistics

For the analysis of microglia cell density and spine density, cells or spines were counted manually in maximum intensity projections of the confocal image stacks using the ‘Cell Counter’ plugin of Fiji image processing package [available at https://fiji.sc/; (Schindelin et al., 2012)].

Spine head volumes were assessed in the confocal image stacks using the Imaris x64 (version 9.5.0) software. The surface tool with the ‘split touching object’ option enabled was used to measure the volume of spine heads. Files were stored as .ims.

Confocal images of PI stained cultures were processed and analyzed using the Fiji image processing package. After background subtraction (rolling ball radius 50 px), images were binarized and PI positive particles were displayed and counted using the ‘Analyze Particles’ function. Values were normalized to the mean value of the control group.

Confocal image stacks of heterozygous *C57BL/6-Tg(TNFa-eGFP)* cultures were processed and analyzed as previously described (Lenz et al., 2020) using the Fiji image processing package. Mean fluorescence intensity of the culture area was normalized to the mean value of fluorescence intensity of the sham-stimulated cultures in the respective round.

Dendritic morphologies were assessed using the Neurolucida^®^ 360 software (version 2020.1.1). Cells were semi-automatically reconstructed with the ‘user guided tree reconstruction’ function. Reconstructions were saved as .DAT files and analysis was performed in the Neurolucida Explorer (version 2019.2.1).

To analyze microglia morphology in *HexB-tdTom* cultures, confocal image stacks were processed and analyzed using the Fiji image processing package. Of each z-stack a maximum intensity projection was generated and binarized using the ‘Trainable Weka Segmentation’ plugin (Arganda-Carreras et al., 2017). The same classifier was applied to all images of the same microglia over the recorded 3 h. After removing outliers (radius = 2 px, threshold = 50), microglia scanning density and microglia domain of each image were manually assessed as previously described (Pfeiffer et al., 2016). Values were normalized to the mean.

For the analysis of microglial morphology in acute mouse cortical slices, stacks of single cells were also processed and analyzed using the Fiji image processing package. First, each stack was processed using the ‘despeckle’ function, then a maximum intensity projection was generated. After removing outliers (radius = 3 px, threshold = 50), the image was binarized as described before. Again, outliers were removed (radius = 2 px, threshold = 50), microglia scanning density and microglia domain of each image were manually assessed.

Single cell recordings were analyzed off-line using Clampfit 11 of the pClamp11 software package (Molecular Devices). sEPSC and mEPSC properties were analyzed using the automated template search tool for event detection (Lenz et al., 2021). Input resistance was calculated for the injection of −100 pA current at a time frame of 200 ms with maximum distance to the Sag-current. Resting membrane potential was calculated as the mean baseline value. AP detection was performed using the input/output curve threshold search event detection tool, and the AP frequency was assessed upon the number of APs detected during the respective injection step. 4 cells in the microglia-depleted (BLZ945-treated) sham stimulated group were excluded in the analysis of intrinsic properties (Figure S4) due to loss of the integrity of the patch during recording of the I-V-curve.

Statistical evaluation of the multi-scale modeling was implemented in R (4.0.3; https://www.R-project.org/) and R Studio (1.3.1093; http://www.rstudio.com/) integrated development environment. We ran generalized linear mixed models (GLMM) with predictors TREATMENT (two levels: control, PLX3397) and COMPARTMENT (three levels: soma, apical and basal dendrites). GLMM allows modeling dependent variables from different distributions and model both fixed and random effects (Stroup, 2012). The null model contained the cell as random intercept, and we added each predictor and their interaction terms one-by-one to the subsequent models. The Bayesian information criterion (BIC) was used to compare the current model with the previous one. We selected the winning model if the ΔBIC was at least 10 units less than for the null or previous model.

Affymetrix GeneChip™ microarray data (CEL files) were analyzed using the Affymetrix Transcriptome Analysis Console (TAC version 4.0.2.15). Gene expression was considered significantly different when FDR p-value < 0.05 and fold change < −2 or > 2. Differentially expressed well-annotated genes were considered ‘microglia-specific’ (Figure 1) if they were part of highly specific microglia markers found by Chiu et al. (Chiu et al., 2013) and ‘microglia-related’ if there was literature found that showed expression of these genes in microglia. A full list of differentially expressed genes including predicted genes is provided in supplementary table 1.

RT-qPCR data were analyzed as previously described (Lenz et al., 2020) with the Bio-Rad CFX Maestro 1.0 software package using the ΔΔCq method with *Gapdh* as reference gene. Values were normalized to the mean value of the respective vehicle-treated control group.

Mesoscale cytokine detection assay was analyzed using the MSD DISCOVERY WORKBENCH 4.0 platform. mRNA/protein level correlations were visualized by a linear regression fit and analyzed using non-parametric Spearman’s correlation coefficients (r).

Data were analyzed using GraphPad Prism 7 (GraphPad software, USA). Statistical comparisons were made using non-parametric tests, since normal distribution could not be assured. For column statistics, Mann-Whitney test (to compare two groups) and Kruskal-Wallis-test followed by Dunn’s multiple comparisons were used. For statistical comparison of XY-plots, we used an RM two-way ANOVA test (repeated measurements/analysis) with Sidak’s multiple comparisons. p-values < 0.05 were considered a significant difference. In the text and figures, values represent mean ± standard error of the mean (s.e.m.). * p < 0.05, ** p < 0.01, *** p < 0.001 and not significant differences are indicated by ‘ns’.

### Digital Illustrations

Figures were prepared using Photoshop graphics software (Adobe, San Jose, CA, USA). Image brightness and contrast were adjusted.

## Results

### Pharmacologic depletion of microglia in organotypic tissue cultures

To determine the role of microglia in r(T)MS-induced synaptic plasticity organotypic tissue cultures were treated with the colony-stimulating factor 1 receptor (CSF-1R) antagonist PLX3397, which readily depletes microglia (Coleman et al., 2020; Elmore et al., 2014). Tissue cultures containing the entorhinal cortex and the hippocampus were exposed to 50 nM PLX3397 for at least 18 days *in vitro*, starting immediately after tissue-culture preparation (Figure 1). A robust and almost complete depletion of microglia (∼96% reduction in cell density) was observed in the three-week old tissue cultures, as demonstrated by immunostaining for the microglial marker Iba1 (Figure 1B-D).

Successful depletion of microglia was also confirmed with RNA microarray analysis (Figure 1E-G). We observed a significant reduction of microglia-specific and microglia-related genes in the group treated with PLX3397 (Figure 1E, F). Specifically, gene sets characteristic of M0-, M1-, and M2-classified microglia (Jurga et al., 2020) were reduced in our tissue cultures (Figure 1G). In turn, no major changes in the expression of genes related to astrocytes, oligodendrocytes, and neurons, i.e., synaptic genes were observed following microglia depletion. These findings show that treatment with PLX3397 results in a robust and specific depletion of microglia in three-week-old organotypic tissue cultures.

### CA1 pyramidal neurons in microglia-depleted tissue cultures do not express rMS-induced synaptic plasticity

Microglia-depleted tissue cultures and non-depleted control cultures were stimulated with a 10 Hz stimulation protocol consisting of 900 pulses at 50% maximum stimulator output using a Magstim Rapid^2^ stimulator equipped with an AirFilm^®^ coil (Figure 2). The distance from the coil and the orientation of the stimulated tissue within the electric field were kept constant in all experiments (Figure 2A, B). CA1 pyramidal neurons were patched and α-amino-3-hydroxy-5-methyl-4-isoxazolepropionic acid (AMPA) receptor-mediated miniature excitatory postsynaptic currents (mEPSCs) were recorded 2–4 h after stimulation (Figure 2C). Sham-stimulated controls were treated the same way except for stimulation. In line with previous work (Lenz et al., 2020; Lenz et al., 2015; Vlachos et al., 2012), 10 Hz rMS induced a robust strengthening of excitatory synapses in CA1 pyramidal neurons of control tissue cultures, as demonstrated by increased mean mEPSC amplitude, half width, area, and frequency (Figure 3A-C). Conversely, no changes in mEPSCs were observed in microglia-depleted tissue cultures following rMS (Figure 3D-E). These results demonstrate that the presence of microglia is required for rMS-induced synaptic plasticity.

### Microglia depletion does not affect cell viability

The neurotrophic and neuroprotective effects of microglia are well recognized in the field (e.g., (Freria et al., 2020; Jin et al., 2017)). Although no obvious signs of cell death were observed in our experiments and despite the fact that baseline mEPSC properties were comparable between the two groups (c.f., Figure 3C and F), we decided to err on the side of caution and tested for alterations in cell viability as a potential confounding factor of rMS-induced synaptic plasticity. In these experiments propidium iodide (PI) staining was used to assess viability and cell death in our preparations [Figure 4; c.f., (Lenz et al., 2020)]. PI is a cell-membrane impermeant fluorescent molecule that binds to DNA. Hence, PI can be used as a marker for membrane integrity when applied to living tissue. Tissue cultures treated with PLX3397 (50 nM) or vehicle-only controls were stained with PI and NMDA treated tissue cultures (50 µM, 4 h) served as a positive control in these experiments (Figure 4A). The PI-signal was comparatively low in three-week-old control cultures and age- and time-matched PLX3397-treated preparations, while a significant increase in PI-signal was detected in the NMDA-treated group (Figure 4A). We conclude from these results that the inability of CA1 pyramidal neurons to express rMS-induced synaptic plasticity is not based on altered cell viability or cell death in the absence of microglia.

**Figure 4:**
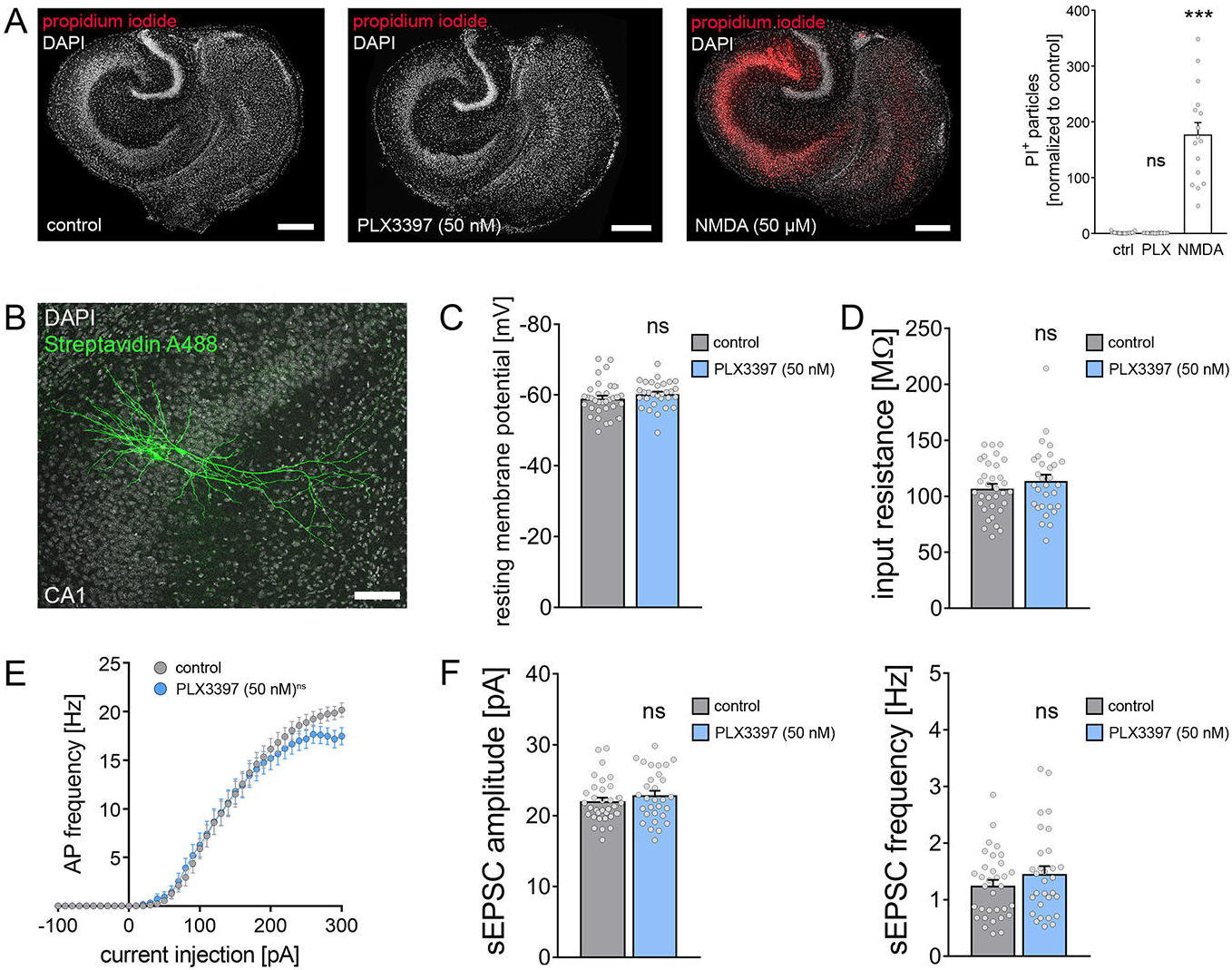
Depletion of microglia does not affect cell viability and basic functional properties of CA1 pyramidal neurons. (A) Tissue cultures stained with propidium iodide [left: vehicle control, middle: PLX3397 (50 nM, 18 d), right: NMDA (50 μM, 4 h)], group data of the quantified propidium iodide signals is shown in the graph on the right (n_control_ = 29 cultures, n_PLX(50nM)_ = 28 cultures, n_NMDA_ = 16 cultures; Kruskal-Wallis-test followed by Dunn’s post-hoc correction). Scale bars, 200 µm. (B) Examples of patched, recorded and posthoc identified CA1 pyramidal neurons. Scale bar, 100 µm. (C, D) Group data of passive membrane properties of CA1 pyramidal neurons in microglia-depleted, i.e., PLX3397 (50 nM, 18 days) treated tissue cultures and control cultures (n_control_ = 33 cells, n_PLX3397_ = 30 cells; Mann-Whitney test). (E) Group data of action potential frequencies of CA1 pyramidal neurons in the respective groups (n_control_ = 33 cells, nPLX3397 = 30 cells, RM two-way ANOVA followed by Sidak’s multiple comparisons). (F) Group data of AMPA receptor-mediated spontaneous excitatory postsynaptic currents (sEPSCs) recorded from CA1 pyramidal neurons revealed no significant changes in excitatory neurotransmission in microglia-depleted tissue cultures (n_control_ = 33 cells, n_PLX3397_ = 30 cells; Mann-Whitney test). Individual data points are indicated by grey dots. Values represent mean ± s.e.m (*** p < 0.001; ns, not significant differences).

### No major functional and structural changes of CA1 pyramidal neurons in microglia-depleted tissue cultures

Neuronal excitability and morphology are expected to have a major impact on the outcome of electromagnetic stimulation (Aberra et al., 2020; Shirinpour et al., 2021). Therefore, we tested for the effect of microglia depletion on structural and functional properties of CA1 pyramidal neurons in our experiments (Figure 4-6). Another set of CA1 pyramidal neurons was recorded in control and microglia-depleted tissue cultures and stained posthoc for morphological analysis (Figure 4B; c.f., Figures 5 and 6).

**Figure 5:**
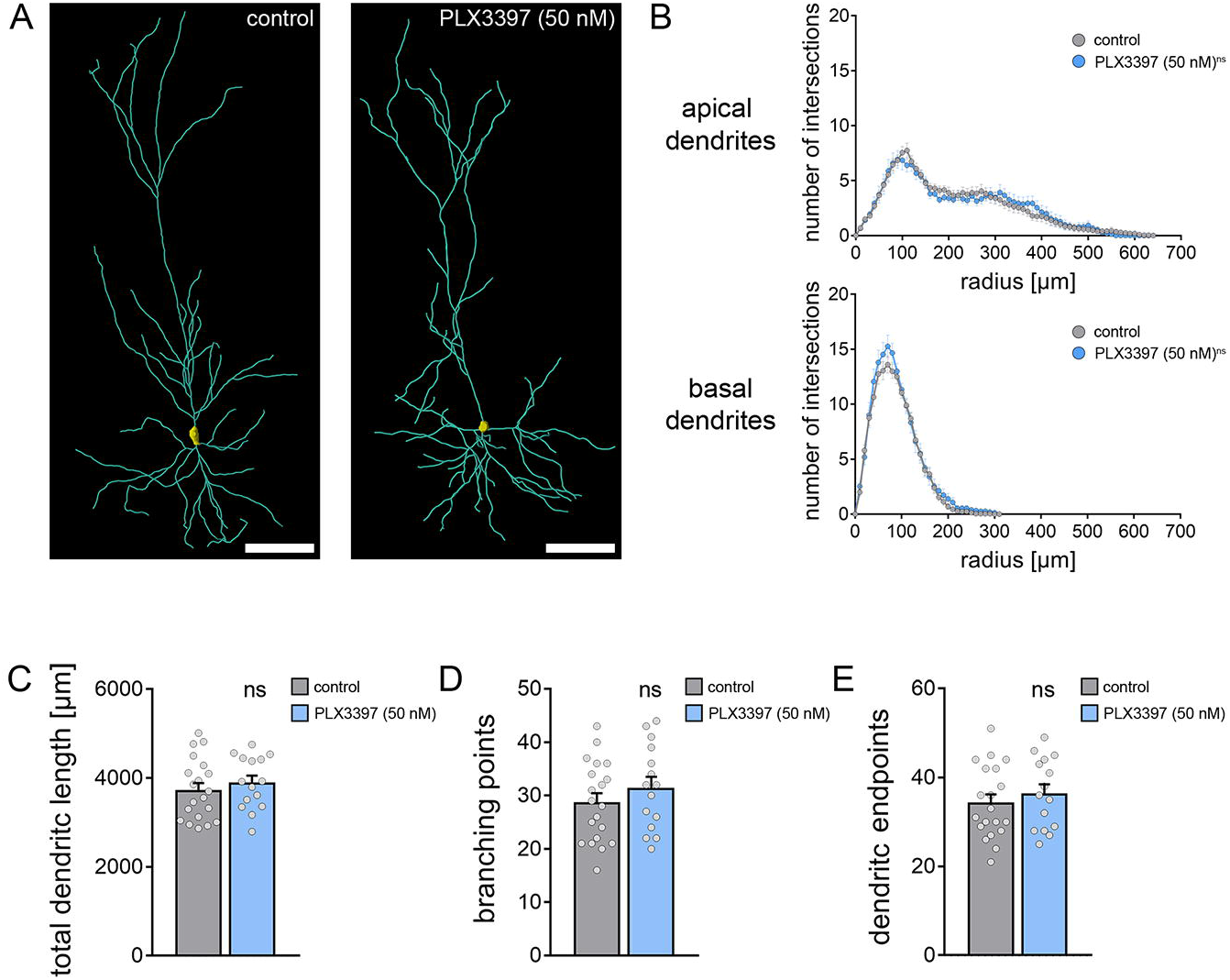
Dendritic morphology of CA1 pyramidal cells is not affected in microglia-depleted tissue cultures. (A) Examples of three dimensionally reconstructed CA1 pyramidal neurons in non-depleted controls and PLX3397 treated, i.e., microglia-depleted tissue cultures. Scale bars, 100 µm. (B) Sholl analysis of apical and basal dendrites from reconstructed CA1 neurons in the respective groups (n_control_ = 20 cells, n_PLX3397_ = 15 cells; RM two-way ANOVA followed by Sidak’s multiple comparisons). (C-E) Group data of additional morphological parameters from the same set of reconstructed CA1 pyramidal neurons in microglia-depleted tissue cultures and vehicle-only treated control cultures (n_control_ = 20 cells, n_PLX3397_ = 15 cells; Mann-Whitney test). Individual data points are indicated by grey dots. Values represent mean ± s.e.m (ns, not significant differences).

**Figure 6:**
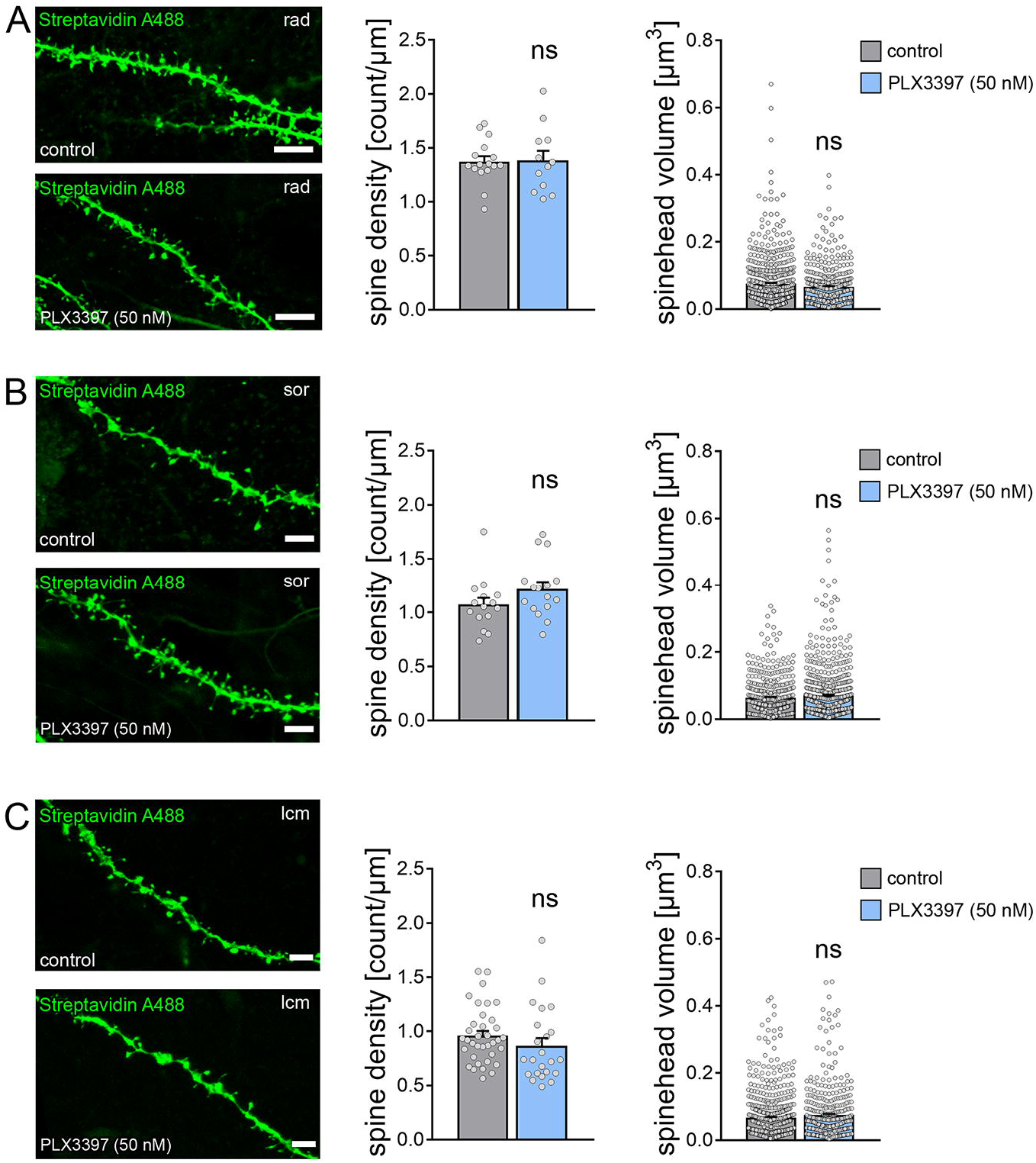
Dendritic spines of CA1 pyramidal neurons are not altered in microglia-depleted tissue cultures. (A-C) Examples of dendritic segments and group data of spine densities and spine volumes from patched and posthoc identified CA1 pyramidal neurons in stratum radiatum (rad, A), stratum oriens (sor, B) and stratum lacunosum-moleculare (lcm, C) of PLX3397 treated, i.e., microglia-depleted tissue cultures and control cultures (rad density: n_control_ = 16 dendritic segments, n_PLX3397_ = 12 dendritic segments; rad volume: n_control_ = 578 spines, n_PLX3397_ = 393 spines; oriens density: n_control_ = 15 dendritic segments, n_PLX3397_ = 16 dendritic segments; oriens volume: n_control_ = 450 spines, n_PLX3397_ = 703 spines; lcm density: n_control_ = 36 dendritic segments, n_PLX3397_ = 23 dendritic segments; lcm volume: n_control_ = 655 spines, n_PLX3397_ = 405 spines; Mann-Whitney test). Scale bars, 3 µm. Individual data points are indicated by grey dots. Values represent mean ± s.e.m (ns, not significant differences).

PLX3397-mediated microglia depletion did not affect the basic functional properties of CA1 pyramidal neurons, i.e., resting membrane potential (Figure 4C) and input resistance (Figure 4D). A slight but non-significant difference in action potential frequency was observed for high current injections between the two groups (Figure 4E). Likewise, amplitudes and frequencies of AMPA-receptor mediated spontaneous EPSC were not significantly different between the groups (Figure 4F). These results indicate that microglia depletion does not affect basic functional properties and synaptic activity of CA1 pyramidal neurons.

Consistent with these findings, Sholl analysis of the reconstructions of apical and basal dendrites did not show any major effects of microglia depletion on the dendritic morphologies of CA1 pyramidal neurons (Figure 5A, B). We did not observe any significant differences in total dendritic length (Figure 5C), the number of branching points (Figure 5D), or the number of endpoints (Figure 5E).

Finally, a detailed morphological analysis of CA1 dendritic spines (Figure 6) confirmed our sEPSC recordings by demonstrating no significant differences in spine densities and spine-head volumes between the two groups in CA1 stratum radiatum (Figure 6A), stratum oriens (Figure 6B) and stratum lacunosum-moleculare (Figure 6C).

Taken together, we conclude that microglia depletion does not cause major structural and functional changes in CA1 pyramidal neurons that could readily explain the inability of neurons to express rMS-induced synaptic plasticity in the absence of microglia.

### Realistic multi-scale computer modeling predicts no major differences in rMS-induced depolarization and intracellular Ca^2+^ levels

We further evaluated the effects of 10 Hz rMS on CA1 pyramidal neurons using multi-scale computational modeling to link the physical input parameters of rMS to dendritic morphologies and subcellular neural effects [Figure 7; (Shirinpour et al., 2021)]. The 3D reconstructed morphologies from recorded CA1 pyramidal neurons in microglia-depleted and non-depleted tissue cultures (c.f., Figure 5) were used in these experiments (Figure 7A). Both membrane voltage and calcium concentrations were assessed in this modeling approach (Figure 7B-F). We estimated the minimum synaptic weight just below the firing threshold of the CA1 pyramidal neurons. Consistent with our experimental data the synaptic weights of the model did not differ in CA1 pyramidal neurons from depleted and non-depleted tissue cultures (Figure 8D).

**Figure 7:**
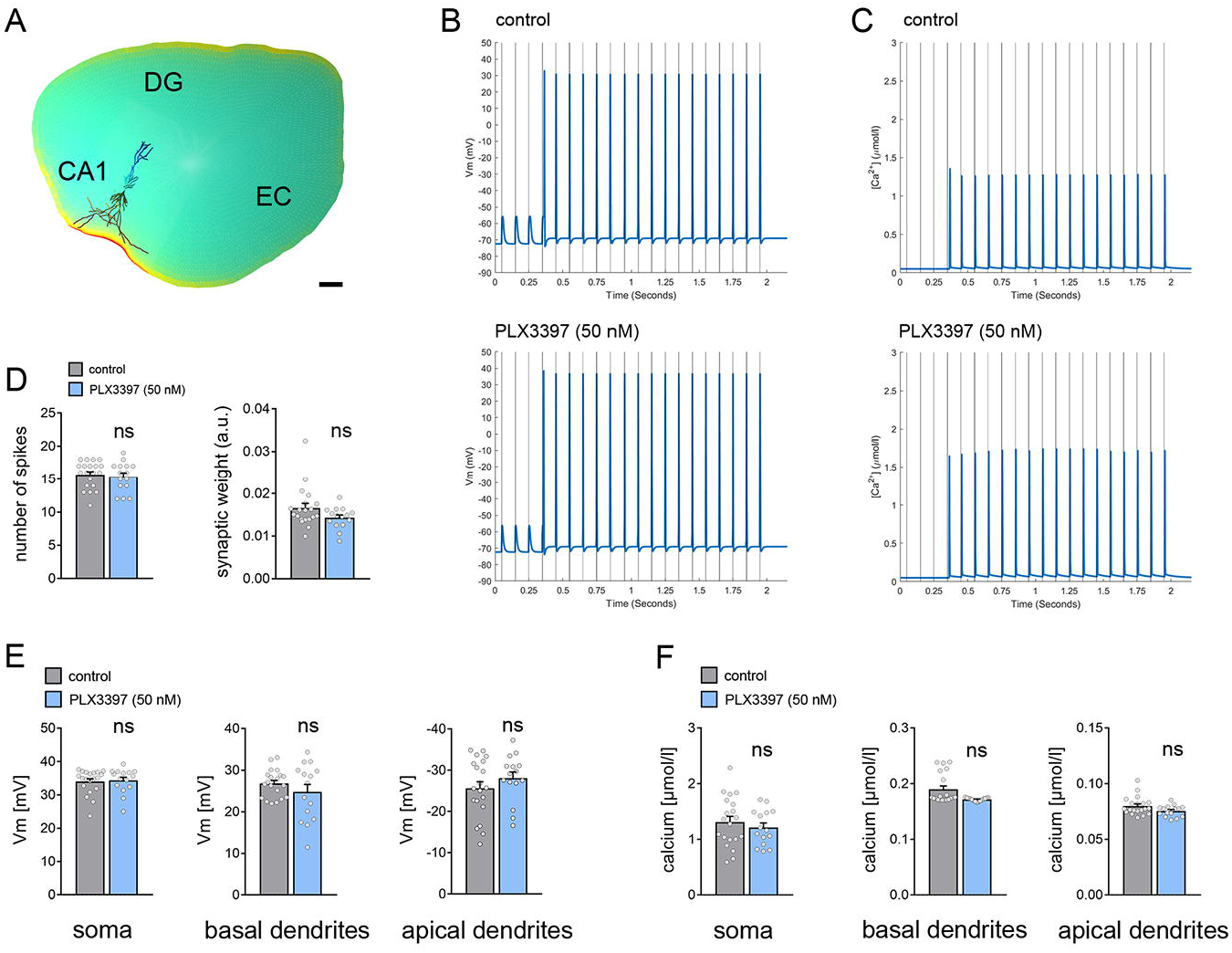
Multi-scale computer modeling of repetitive magnetic stimulation (rMS) (A) Neuronal responses to rMS were modeled in realistic dendritic morphologies from reconstructed CA1 pyramidal neurons in a stereotypic tissue culture environment. EC, entorhinal cortex; DG, dentate gyrus. Scale bar, 200 µm. (B, C) Changes in membrane voltage (Vm; B) and intracellular calcium levels (Ca^2+^; C) were assessed for a train of 20 pulses at 10 Hz at the constant stimulation intensity used in the experimental setting. CA1 pyramidal neurons in both conditions showed a delayed suprathreshold response upon repetitive magnetic stimulation. (D) Both the number of cellular spikes upon stimulation and the synaptic weight did not show a significant difference between CA1 pyramidal neurons from microglia-depleted and non-depleted control cultures (n_vehicle-only_ = 20 cells, n_PLX3397_ = 15 cells; GLMM). (E) In the same set of cells the peak membrane voltage difference in response to magnetic stimulation was modeled in the somatic, apical and basal dendritic compartments. No differences were observed between the two groups, respectively. (F) Analysis of stimulus-triggered changes in intracellular calcium levels. No changes in both the dendritic and the somatic compartments were evident between CA1 pyramidal neurons of microglia-depleted tissue cultures and control cultures. Individual data points are indicated by grey dots. Values represent mean ± s.e.m. (ns, not significant difference).

**Figure 8:**
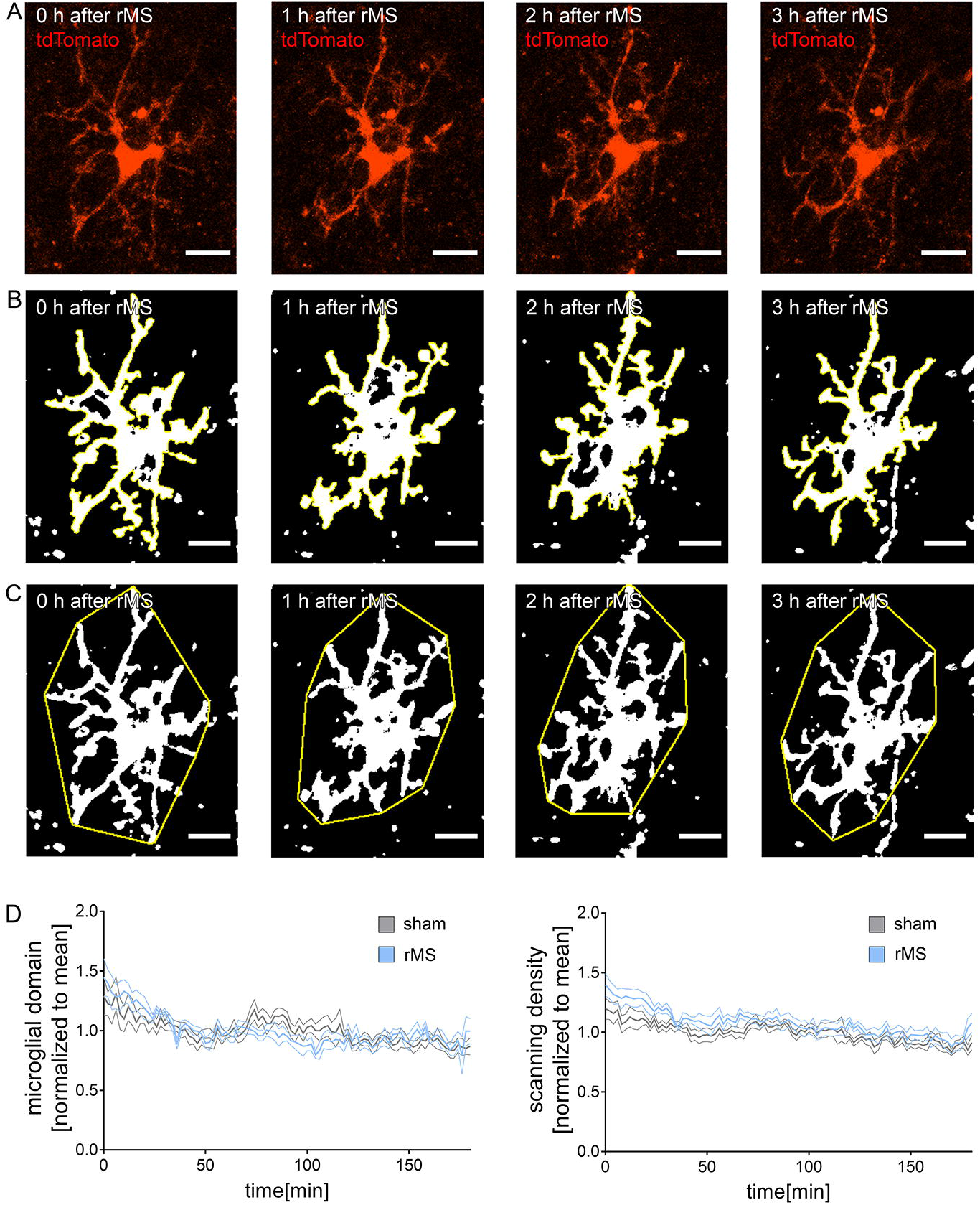
Repetitive magnetic stimulation (rMS) does not affect microglia morphology. (A-C) Examples of tdTomato expressing microglia in hippocampal area CA1 imaged from *HexB^tdT/tdT^*cultures over a period of three hours following 10 Hz rMS (2 min intervals). Maximum intensity projections of image-stacks (A), microglial scanning densities (B), and microglial domains (C) are illustrated. Scale bars, 15 µm. (D) Group data of microglial domains and scanning densities from rMS-stimulated and sham-stimulated tissue cultures (n_sham_ = 6 cells, n_rMS_ = 6 cells from 6 cultures in each group; RM two-way ANOVA followed by Sidak’s multiple comparisons). Values represent mean ± s.e.m. (ns, not significant difference).

We compared the peak spike values for the membrane voltage. The winning model (ΔBIC = 341.24; F_2,102_ = 1387.59, p = 2.2 × 10^-16^) employed the compartment as predictor. Confirming our expectations, the analysis revealed significantly weaker peak values in the apical (t = −48.524, p = 2.2 × 10^-16^) and basal dendrites (t = −6.563, p = 2.23 × 10^-9^). However, the treatment had no effect on either the number of spikes (Figure 7D) or the peak action potential values in any compartment (Figure 7E).

We continued by comparing the peak calcium-concentration values extracted from the compartments and treatment conditions. As for the voltage data, the winning model (ΔBIC = 189.3875; F_2,102_ = 283.51, p = 2.2 × 10^-16^) utilized the compartment as predictor. Again, the analysis revealed significantly weaker peak values in the apical (t = −21.64, p = 2.2 × 10^-16^) and basal dendrites (t = −19.75, p = 2.2 × 10^-16^). However, the treatment had no major effects on the peak calcium-concentration levels in compartments investigated (Figure 7F).

### Repetitive magnetic stimulation does not affect structural properties of microglia

After demonstrating that structural and functional differences of CA1 pyramidal neurons do not explain our major findings, i.e., the inability of neurons to express rMS-induced synaptic plasticity in the absence of microglia, we considered the possible effects of rMS on the structural and functional properties of microglia (Figures 8 and 9).

**Figure 9:**
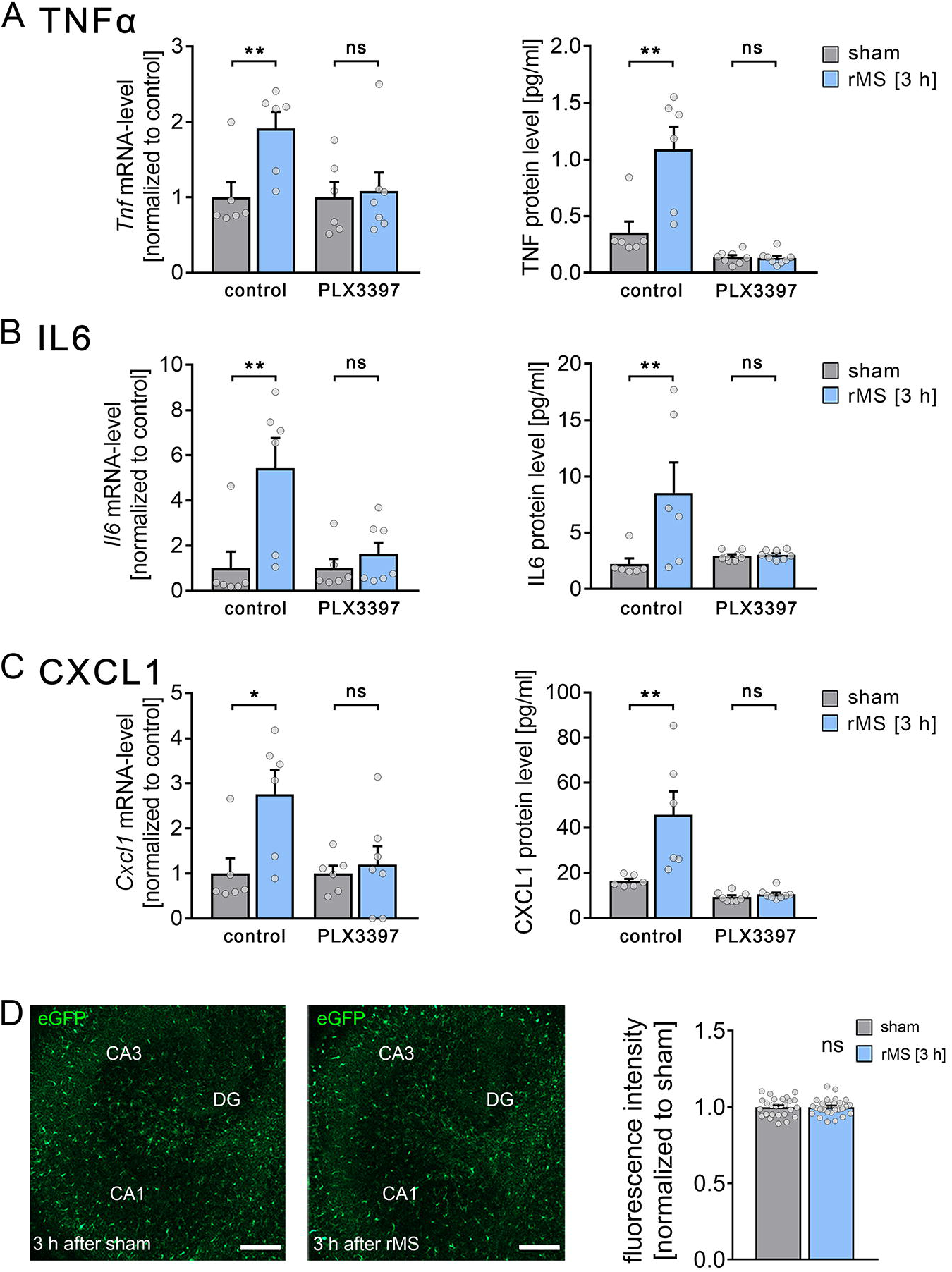
Repetitive magnetic stimulation (rMS) triggers the expression and release of plasticity-promoting microglial factors. (A-C) Group data of mRNA and protein levels of (A) tumor necrosis factor alpha (TNFα), (B) interleukin 6 (IL6) and (C) chemokine ligand 1 (CXCL1) in cultures and microglia-depleted cultures (PLX3397 treated cultures) after 10 Hz rMS-stimulated tissue cultures and sham-stimulated controls. (n_control_ = 6 cultures or culturing medium samples respectively for each experimental condition, n_PLX3397 mRNA sham_ = 6 cultures, n_PLX3397 mRNA rMS_ = 7 cultures; n_PLX3397 protein levels_ = 8 culturing medium samples respectively for each experimental condition; Mann-Whitney test, Ucontrol Tnf mRNA = 2, U_control TNF protein_ = 2, U_control Il6 mRNA_ = 2, U_control IL6 protein_ = 2, U_control Cxcl1 mRNA_ = 3, U_control CXCL1 protein_ = 0). (D) Sample images and group data of eGFP fluorescence intensity of rMS-stimulated and sham-stimulated tissue cultures prepared from TNF-reporter mice [*C57BL/6-Tg(TNFa-eGFP)*] imaged 3 h after 10 Hz rMS (DG, dentate gyrus; n_sham_ = 26 cultures, n_rMS_ = 27 cultures; Mann-Whitney test). Scale bars, 200 µm. Individual data points are indicated by grey dots. Values represent mean ± s.e.m. (* p < 0.05, ** p < 0.01; ns, not significant difference).

To test for the effects of 10 Hz rMS on microglia morphology and dynamics, tissue cultures were prepared from the recently established transgenic *HexB-tdTom* mouse line, which expresses the red fluorescent protein tdTomato under the control of the Hexoaminidase B promotor (HexB) (Masuda et al., 2020). Of all tdTomato expressing cells, 97% also showed a positive Iba1 staining, while in 98% of all Iba1 positive cells, tdTomato signal was also detected (Figure S1). Apparently, the vast majority of microglia are readily identified in three-week-old tissue cultures prepared from *HexB-tdTom* mice.

We next employed live-cell microscopy and imaged microglia, i.e., tdTomato expressing cells in CA1 stratum radiatum. Confocal image stacks were obtained every 2 minutes for 3 hours immediately after 10 Hz rMS or sham stimulation (Figure 8). In these experiments, we did not observe any major rMS-related changes in microglia morphology. A detailed analysis of microglia dynamics in maximum-intensity projections (Figure 8A-C) revealed no significant changes in dynamic microglial domains and scanning densities among rMS-stimulated and sham-stimulated tissue cultures (Figure 8D). We conclude that 10 Hz rMS does not lead to major changes in microglia morphology and dynamics.

### Repetitive magnetic stimulation triggers a release of plasticity-promoting cytokines from microglia

We then considered whether 10 Hz rMS triggers an activation of microglia in the absence of major changes in microglia morphology. The plasticity-modulating effects of cytokines secreted by activated microglia are well-established (e.g., Cao et al., 2014; Chai et al., 2019; Habbas et al., 2015; Heir and Stellwagen, 2020; Lenz et al., 2020; Maggio and Vlachos, 2018; Santello et al., 2011; Stellwagen and Malenka, 2006; Tancredi et al., 2000). Specifically, TNFα and IL6 have been linked to the ability of neurons to express synaptic plasticity (Lenz et al., 2015; Rizzo et al., 2018; Stellwagen and Malenka, 2006; Tancredi et al., 2000). In this context, we were recently able to demonstrate that low concentrations of TNFα promote the ability of neurons to express synaptic plasticity (Maggio and Vlachos, 2018). Therefore, we theorized that 10 Hz rMS could trigger the production and/or secretion of TNFα and other cytokines at low, i.e., plasticity-promoting concentrations.

We performed qPCR-analyses and protein-detection assays in another set of 10 Hz stimulated and sham-stimulated microglia-depleted and non-depleted tissue cultures (Figure 9A-C and Figure S2). Indeed, 10 Hz rMS triggered an increase in TNFα at both the mRNA and protein levels that was not evident in microglia-depleted tissue cultures (Figure 9A). Notably, reduced baseline levels of TNFα protein were observed in the microglia-depleted tissue cultures, which is consistent with the suggestion that microglia are a major source of TNFα in our preparations (n_control, non-depleted_ = 6 cultures, n_control, PLX3397_ = 8 cultures; Mann-Whitney test). Moreover, a considerable increase in IL6 and CXCL1 was detected only in non-depleted tissue cultures (Figure 9B, C). Notably, IL1β could only be detected at the lower detection limit and did not increase significantly upon stimulation.

We further excluded an rMS-induced pathological activation of microglia in tissue cultures prepared from a transgenic TNFα reporter mouse line, which expresses the enhanced green fluorescent protein under the control of the TNF-promoter, *C57BL/6-Tg(TNFa-eGFP).* In a previous study, we found a strong pathological activation of microglia, i.e., strong increase in eGFP-expression, in the presence of bacterial lipopolysaccharides (Lenz et al., 2020). Live-cell microscopy revealed no obvious changes in eGFP fluorescence following 10 Hz rMS (Figure 9D), which is in line with a moderate production and release of cytokines (Figure 9A-C) in the absence of obvious morphological changes of microglia (Figure 8). Taken together, these results suggest that 10 Hz rMS induces a physiological increase and release in microglial cytokines, e.g., TNFα and IL6.

### Substitution of plasticity-promoting cytokines during stimulation rescues rMS-induced synaptic plasticity in microglia-depleted tissue cultures

We then tested whether the exogenous substitution of TNFα and IL6 rescues the ability of CA1 pyramidal neurons to express rMS-induced synaptic plasticity in the absence of microglia (Figure 10). TNFα (5 ng/ml) and IL6 (2.5 ng/ml) were added to the medium during stimulation only, and AMPA-receptor mediated mEPSCs were again recorded from CA1 pyramidal neurons of microglia-depleted tissue cultures between 2-4 h after stimulation (Figure 10A).

**Figure 10:**
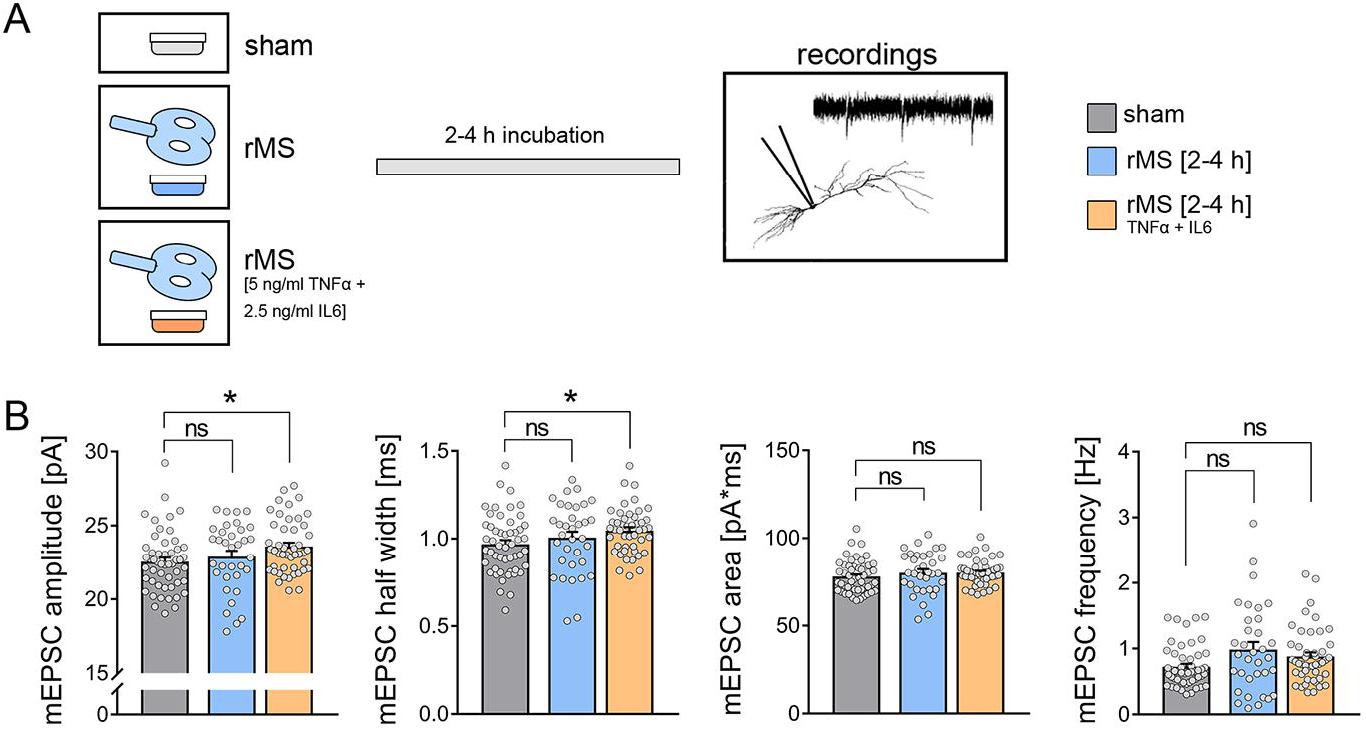
Substitution of pro-inflammatory cytokines during stimulation in microglia-depleted tissue cultures rescues rMS-induced synaptic plasticity. (A) Microglia depleted cultures were stimulated with 10 Hz rMS or sham stimulation. To simulate the rMS-induced cytokine release on tissue level, the rMS-stimulation of a third set of cultures was performed in the presence of TNFα (5 ng/ml) and IL6 (2.5 ng/ml). (B) Group data of AMPA receptor-mediated miniature excitatory postsynaptic currents (mEPSCs) recorded from CA1 pyramidal neurons in sham-stimulated and 10 Hz rMS-stimulated cultures (n_sham_ = 51 cells, n_rMS_ = 34 cells, n_rMS-TNF+IL6_ = 46 cells; Kruskal-Wallis-test followed by Dunn’s post-hoc correction). Individual data points are indicated by grey dots. Values represent mean ± s.e.m (* p < 0.05; ns, not significant differences).

Indeed, in these experiments rMS-induced synaptic strengthening, i.e., a significant increase in mean mEPSC amplitude and half width was observed (Figure 10B). Of note, exposure to TNFα and IL6 at the same concentration and duration did not affect mEPSC amplitudes; only mEPSC frequencies were increased (n_sham_ = 20 cells, n_sham+cytokines_ = 22 cells; mEPSC amplitude: 24.27 ± 0.56 pA (sham) vs. 24.86 ± 0.43 pA (sham+cytokines), ns; mEPSC frequency: 0.81 ± 0.06 Hz (sham) vs. 0.95 ± 0.05 Hz (sham+cytokines), p < 0.05*; Mann-Whitney test). We conclude that TNFα and IL6 released from microglia during stimulation is required for rMS-induced synaptic potentiation.

### *In vivo* microglia depletion prevents rTMS-induced excitatory synaptic plasticity in the mouse mPFC

Finally, we tested whether the presence of microglia is required for rTMS-induced plasticity of synaptic transmission *in vivo* (Figure 12; Kloosterboer and Funke, 2019; Lenz et al., 2016; Thimm and Funke, 2015). *In vivo* microglia depletion was achieved through daily application of the CSF-1R inhibitor BLZ945 (200 mg kg^-1^, oral gavage, for 7 consecutive days prior to experimental assessment; Hagemeyer et al., 2017; Masuda et al., 2020). Anesthetized microglia-depleted and non-depleted mice were stimulated with the same 10 Hz stimulation protocol as in our *in vitro* experiments. Two hours after the stimulation, frontal brain sections containing the medial prefrontal cortex were prepared and AMPA-receptor meditated sEPSCs were recorded from superficial pyramidal neurons of the mPFC (3-5 hours post stimulation; Figure 11A). BLZ945-induced microglia depletion was confirmed by Iba1 stainings of frontal brain sections (∼90% reduction in cell density in the BLZ945-treated animals; Figure 11B). While *in vivo* microglia-depletion had no significant effects on baseline synaptic transmission, a significant increase in sEPSC frequencies was not observed in the stimulated microglia-depleted animals. Notably, the mean sEPSC amplitude remained unaltered in microglia-depleted and non-depleted animals (Figure 11C). Amplitude-frequency plots confirmed that rTMS promoted an amplitude-independent increase in sEPSC frequencies which was absent - if not reversed - in the mPFC of microglia-depleted animals (Figure 11C, D). These changes occurred in the absence of major changes in active and passive membrane properties in microglia-depleted and non-depleted animals (Figure S3). Similar to our *in vitro* experiments in hippocampal area CA1, no rTMS-induced changes in microglia morphologies were observed in the mPFC of anesthetized mice (Figure S4). These results demonstrate the essential, i.e., plasticity-promoting, role of microglia for rTMS-induced changes in network activity and neurotransmission.

**Figure 11:**
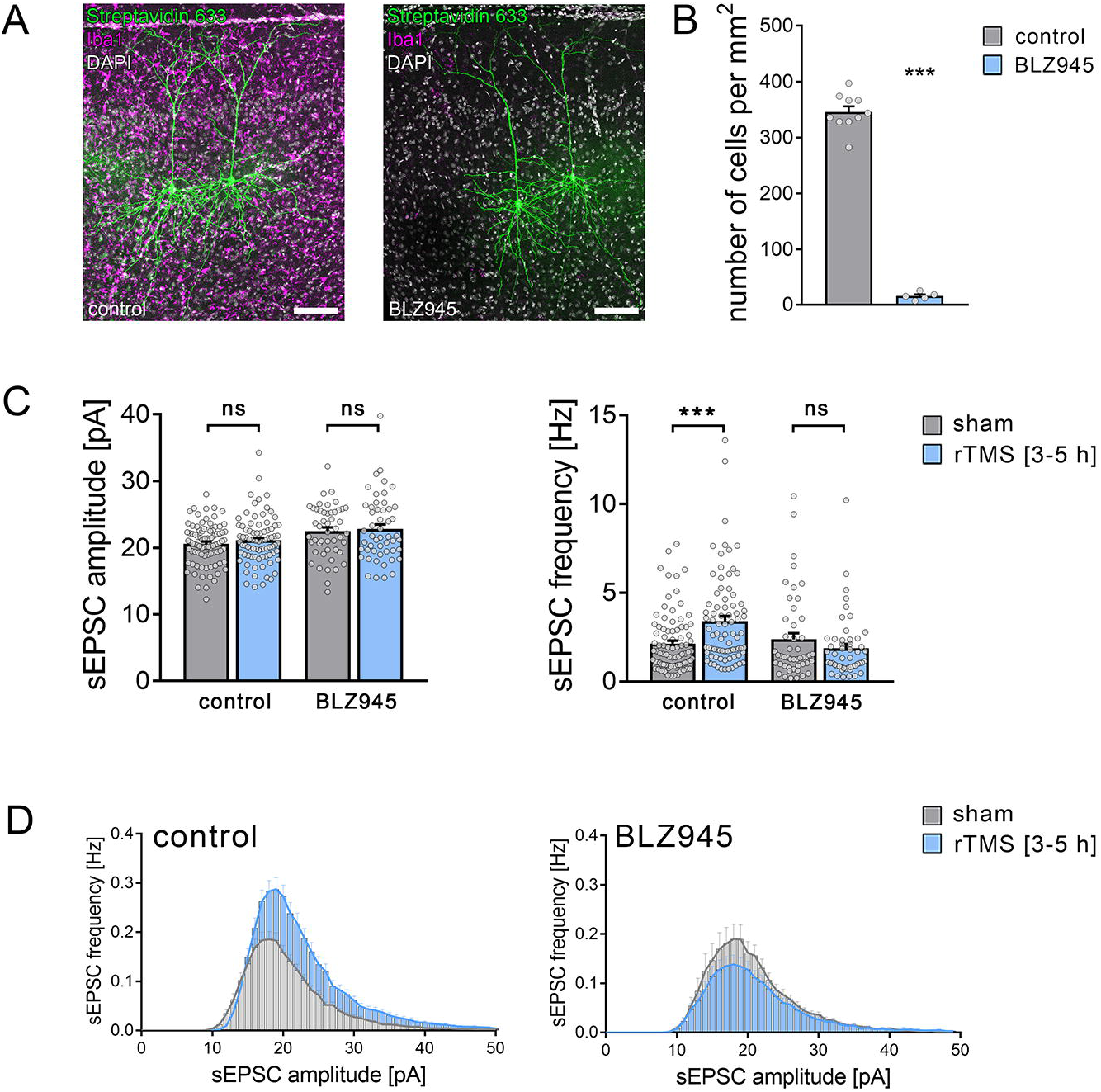
Microglia depletion *in vivo* prevents rTMS-induced synaptic plasticity of superficial pyramidal neurons in the medial prefrontal cortex of adult mice. (A) Post-hoc visualization of superficial pyramidal neurons by streptavidin staining in the medial prefrontal cortex of adult mice. Microglia was visualized by Iba1 immunolabeling. Electrophysiological assessment of excitatory synaptic transmission was performed in frontal sections of non-depleted (left image) and microglia-depleted (BLZ945-treated, right image) animals. Scale bars, 100 µm. (B) Quantification of microglial density confirmed a ∼90% reduction of microglia in the medial prefrontal cortex of BLZ945-treated animals (n_control_ = 10 animals, n_microglia-depleted (BLZ945)_ = 5 animals; Mann-Whitney test. U = 0) (C) 10 Hz repetitive transcranial magnetic stimulation (rTMS) or sham stimulation were performed in anesthetized adult mice and tissue sections were prepared 2 hours after stimulation. After slice recovery, excitatory synaptic transmission recordings were performed 3-5 hours post stimulation. In sections from non-depleted animals, a significant increase in the sEPSC frequency could be detected while die sEPSC amplitude was unchanged. Notably, no changes in excitatory neurotransmission were detected in microglia-depleted tissue sections (non-depleted animals: n_sham_ = 89 cells from 6 animals, nrTMS = 81 cells from 6 animals; microglia-depleted (BLZ945-treated animals): n_sham_ = 48 cells from 2 animals, n_rTMS_ = 51 cells from 3 animals; Mann-Whitney test. U_sEPSC frequency, non-depleted_ = 2324). (D) sEPSC amplitude-frequency-plots from superficial pyramidal cells in the mPFC of 10 Hz-rTMS or sham-treated animals (non-depleted or microglia-depleted). While excitatory synaptic plasticity was evident in non-depleted sections, no changes were detected after microglia-depletion (RM two-way ANOVA following Sidak’s multiple comparisons test). Individual data points are indicated by grey dots. Values represent mean ± s.e.m (*** p < 0.001; ns, not significant differences).

## Discussion

The experiments of this study demonstrate that the presence of microglia is required for the induction of synaptic plasticity triggered by 10 Hz rTMS in brain tissue of mice (*in vitro* and *in vivo*). In this context, a robust functional activation of microglia was observed as reflected by the gene expression and release of TNFα and IL6 (but not IL1b). Indeed, substitution of TNFα and IL6 rescued the ability of neurons to express synaptic plasticity in microglia-depleted brain tissue cultures. The changes occurred in the absence of major morphological and functional changes of the stimulated neurons in the microglia-depleted brain tissue, and no obvious changes of microglia morphologies and their dynamics were observed after 10 Hz electromagnetic stimulation. These results demonstrate that rTMS effectively modulates microglia function, i.e., cytokine release, in stimulated brain regions, thereby identifying a novel mechanism of action which may explain some of the beneficial effects of rTMS seen in healthy individuals and in patients.

During recent years experimental evidence has suggested that rTMS-induced after-effects are mediated by “LTP-like” plasticity mechanisms (Ziemann, 2004). This evidence is based on physiological characteristics and pharmacological analogies between studies performed at the system level in human subjects (Brown et al., 2020; Korchounov and Ziemann, 2011; Ziemann et al., 2015) and data obtained from animal models (Hoogendam et al., 2010; Vlachos et al., 2012). Specifically, the effects of pharmacological modulation of NMDA receptors have been interpreted as evidence for “LTP-like” plasticity, considering the relevance of NMDA receptors in LTP-induction (Brown et al., 2020; Lenz et al., 2015; Vlachos et al., 2012). Our own past work demonstrated that pharmacological inhibition of NMDA receptors or L-Type voltage-gated calcium channels block the ability of neurons to express synaptic plasticity induced by 10 Hz rMS (Lenz et al., 2015; Vlachos et al., 2012). Consistent with these findings, stimulation in Ca^2+^-free external solution failed to induce synaptic plasticity, thus confirming the relevance of Ca^2+^-dependent signaling pathways in rMS-induced synaptic plasticity (Lenz et al., 2015). Notably, NMDA receptors and L-Type voltage-gated calcium channels are also expressed on microglia (Hopp, 2021; Murugan et al., 2011). In fact, robust experimental evidence exists demonstrating that the modulation of intracellular Ca^2+^ levels regulate important microglia functions (Laprell et al., 2021). Hence, some of the results obtained in animal models and human studies could be explained—at least in part—by differential regulation of calcium signaling in microglia and neurons. In this context, additional research is required to clarify whether rTMS acts on microglia directly or whether microglia are activated indirectly by sensing rTMS-induced changes in neuronal activity, e.g., via microglial glutamate receptors, purinergic receptors, or other signaling pathways (Cserep et al., 2020).

Regardless of these considerations, the present study demonstrates that the presence of microglia is required for rTMS-induced synaptic plasticity: CA1 pyramidal neurons in microglia-depleted tissue cultures as well as superficial pyramidal neurons in the mPFC of adult mice did not express rTMS-induced excitatory synaptic plasticity. These findings support the recently emphasized impact of glial cells on the neural effects of non-invasive brain stimulation (Gellner et al., 2021). Interestingly, we did not observe any obvious signs of functional and structural alterations of CA1 pyramidal neurons in microglia-depleted tissue cultures. Consistent with these findings, multi-scale compartmental modeling confirmed that basic morphological and functional properties of neurons do not explain the inability of CA1 pyramidal neurons to express rMS-induced plasticity. We must concede, however, that detailed morphological reconstructions of axons were not obtained, as it is difficult to reliably visualize and reconstruct the entire axon morphology of individual neurons. Indeed, computational studies emphasize the relevance of axons and myelination in rTMS-induced synaptic plasticity (Aberra et al., 2020; Fields, 2005; Shirinpour et al., 2021; Wang et al., 2018). It should be noted, though, that we did not observe any differences in network activity in our experimental setting, as neither sEPSC amplitudes nor frequencies were affected in microglia-depleted tissue cultures. Consistent with these findings we did not observe any differences in dendritic spine counts, and no evidence for alterations in oligodendrocyte markers was obtained in our RNA microarray analysis of microglia-depleted tissue cultures. It is thus unlikely that major changes in the structural and functional properties of CA1 pyramidal neurons explain our results, especially considering our rescue experiments in which only short exposure to TNFα and IL6 during stimulation was sufficient to restore rMS-induced synaptic plasticity in the absence of microglia.

10 Hz rMS induced a robust increase in immune-mediator gene expression and TNFα and IL6 release in organotypic tissue cultures, suggesting that rMS activates microglia. Previous studies have demonstrated that proinflammatory cytokines influence excitatory neurotransmission (Cao et al., 2014; Maggio and Vlachos, 2018; Riazi et al., 2015; Sheppard et al., 2019; Stellwagen and Malenka, 2006; Strehl et al., 2014). Specifically, TNFα and IL6 have been implicated as important secreted factors that modulate synaptic transmission (Furukawa and Mattson, 1998; Garcia-Oscos et al., 2012; Stellwagen et al., 2005) and plasticity (Heir and Stellwagen, 2020). In this context it has been shown that TNFα acts as a permissive factor (Becker et al., 2013; Steinmetz and Turrigiano, 2010), where TNFα *per se* does not trigger major changes in postsynaptic strength; rather, it modulates the ability of neurons to express plasticity without affecting baseline synaptic transmission; besides changes in presynaptic glutamate release (Santello et al., 2011) as also reflected by the increased mEPSC frequencies seen in our experiments, in which microglia-depleted tissue cultures were exposed to TNFα and IL6 without stimulation. Indeed, experiments employing classic tetanic electric stimulation showed that low concentrations of TNFα rapidly promote LTP-induction, while high concentrations of TNFα impede the ability of neurons to express LTP (Maggio and Vlachos, 2018). These findings are consistent with the metaplastic effects of TNFα. In line with this suggestion, pathological activation of microglia with bacterial lipopolysaccharides, which triggers strong TNFα production, i.e., ∼10–15 fold increase in TNFα-mRNA and ∼2000 fold increase in TNFα protein levels, occludes the ability of CA1 pyramidal neurons to express 10 Hz rMS-induced synaptic plasticity (Lenz et al., 2020). Together with the results of the present study, these findings demonstrate that microglia have an important role in rTMS-induced synaptic plasticity. They call for a systematic assessment of rTMS-induced microglia plasticity, and raise the intriguing possibility that rTMS recruits metaplasticity by activating microglia (directly or indirectly). Hence, some of the beneficial effects of rTMS seen in patients may reside—at least in part—in the effects of rTMS on microglia function, which also seem to be involved in promoting the ability of neurons to express “LTP-like” plasticity shortly after stimulation.

## DECLARATIONS

### Acknowledgements

We thank Simone Zenker for skillful technical assistance and the Next Generation Sequencing Core Facility (NGS-CF, Faculty of Medicine, University of Freiburg) for assistance in transcriptome analysis. The authors also acknowledge the support by the state of Baden-Württemberg through the possibility of using high performance computing resources (bwHPC; bwUniCluster 2.0).

### Funding

This work was supported by MOTI-VATE graduate school, Faculty of Medicine, University of Freiburg (to AE), the EQUIP Medical Scientist Programme, Faculty of Medicine, University of Freiburg (to ML), Faculty of Medicine of the University of Freiburg (TUR217/21; to ZT), and Deutsche Forschungsgemeinschaft (DFG; Project-ID 259373024 B14–CRC/TRR 167; to AV).

### Author Contributions

Author contributions have been assigned according to CRediT taxonomy. AE: Validation, Formal Analysis, Investigation, Writing-original draft preparation, Visualization, Funding acquisition. DK: Investigation. ZT: Investigation, Formal Analysis. MF: Investigation. MK: Investigation, Formal Analysis. DP: Resources. TM: Resources. MP: Resources. ML: Conceptualization, Methodology, Validation, Formal Analysis, Investigation, Writing-original draft preparation, Visualization, Supervision, Project administration, Funding acquisition. AV: Conceptualization, Methodology, Resources, Writing-original draft preparation, Supervision, Project administration, Funding acquisition.

**Declarations of interest:** none.

## Notes

### Competing Interest Statement

The authors have declared no competing interest.

### Summary of Updates

In the revised manuscript, current data sets have been extended and in vivo rTMS experiments in anesthetized mice have been added.

## References

Aberra, A.S., Wang, B., Grill, W.M., and Peterchev, A.V. (2020). Simulation of transcranial magnetic stimulation in head model with morphologically-realistic cortical neurons. Brain Stimul 13, 175–189.

Akiyoshi, R., Wake, H., Kato, D., Horiuchi, H., Ono, R., Ikegami, A., Haruwaka, K., Omori, T., Tachibana, Y., Moorhouse, A.J., et al. (2018). Microglia Enhance Synapse Activity to Promote Local Network Synchronization. eNeuro 5.

Arganda-Carreras, I., Kaynig, V., Rueden, C., Eliceiri, K.W., Schindelin, J., Cardona, A., and Sebastian Seung, H. (2017). Trainable Weka Segmentation: a machine learning tool for microscopy pixel classification. Bioinformatics 33, 2424–2426.

Badimon, A., Strasburger, H.J., Ayata, P., Chen, X., Nair, A., Ikegami, A., Hwang, P., Chan, A.T., Graves, S.M., Uweru, J.O., et al. (2020). Negative feedback control of neuronal activity by microglia. Nature 586, 417–423.

Becker, D., Zahn, N., Deller, T., and Vlachos, A. (2013). Tumor necrosis factor alpha maintains denervation-induced homeostatic synaptic plasticity of mouse dentate granule cells. Front Cell Neurosci 7, 257.

Brown, J.C., DeVries, W.H., Korte, J.E., Sahlem, G.L., Bonilha, L., Short, E.B., and George, M.S. (2020). NMDA receptor partial agonist, d-cycloserine, enhances 10 Hz rTMS-induced motor plasticity, suggesting long-term potentiation (LTP) as underlying mechanism. Brain Stimul 13, 530–532.

Cao, D.L., Zhang, Z.J., Xie, R.G., Jiang, B.C., Ji, R.R., and Gao, Y.J. (2014). Chemokine CXCL1 enhances inflammatory pain and increases NMDA receptor activity and COX-2 expression in spinal cord neurons via activation of CXCR2. Exp Neurol 261, 328–336.

Chai, H.H., Fu, X.C., Ma, L., Sun, H.T., Chen, G.Z., Song, M.Y., Chen, W.X., Chen, Y.S., Tan, M.X., Guo, Y.W., et al. (2019). The chemokine CXCL1 and its receptor CXCR2 contribute to chronic stress-induced depression in mice. FASEB J 33, 8853–8864.

Chiu, I.M., Morimoto, E.T., Goodarzi, H., Liao, J.T., O’Keeffe, S., Phatnani, H.P., Muratet, M., Carroll, M.C., Levy, S., Tavazoie, S., et al. (2013). A neurodegeneration-specific gene-expression signature of acutely isolated microglia from an amyotrophic lateral sclerosis mouse model. Cell Rep 4, 385–401.

Cirillo, G., Di Pino, G., Capone, F., Ranieri, F., Florio, L., Todisco, V., Tedeschi, G., Funke, K., and Di Lazzaro, V. (2017). Neurobiological after-effects of non-invasive brain stimulation. Brain Stimul 10, 1–18.

Clarke, D., Penrose, M.A., Harvey, A.R., Rodger, J., and Bates, K.A. (2017). Low intensity rTMS has sex-dependent effects on the local response of glia following a penetrating cortical stab injury. Exp Neurol 295, 233–242.

Coleman, L.G., Jr., Zou, J., and Crews, F.T. (2020). Microglial depletion and repopulation in brain slice culture normalizes sensitized proinflammatory signaling. J Neuroinflammation 17, 27.

Cserep, C., Posfai, B., Lenart, N., Fekete, R., Laszlo, Z.I., Lele, Z., Orsolits, B., Molnar, G., Heindl, S., Schwarcz, A.D., et al. (2020). Microglia monitor and protect neuronal function through specialized somatic purinergic junctions. Science 367, 528–537.

Cullen, C.L., and Young, K.M. (2016). How Does Transcranial Magnetic Stimulation Influence Glial Cells in the Central Nervous System? Front Neural Circuits 10, 26.

Elmore, M.R., Najafi, A.R., Koike, M.A., Dagher, N.N., Spangenberg, E.E., Rice, R.A., Kitazawa, M., Matusow, B., Nguyen, H., West, B.L., et al. (2014). Colony-stimulating factor 1 receptor signaling is necessary for microglia viability, unmasking a microglia progenitor cell in the adult brain. Neuron 82, 380–397.

Fields, R.D. (2005). Myelination: an overlooked mechanism of synaptic plasticity? Neuroscientist 11, 528–531.

Forster, E., Zhao, S., and Frotscher, M. (2006). Laminating the hippocampus. Nat Rev Neurosci 7, 259–267.

Freria, C.M., Brennan, F.H., Sweet, D.R., Guan, Z., Hall, J.C., Kigerl, K.A., Nemeth, D.P., Liu, X., Lacroix, S., Quan, N., et al. (2020). Serial Systemic Injections of Endotoxin (LPS) Elicit Neuroprotective Spinal Cord Microglia through IL-1-Dependent Cross Talk with Endothelial Cells. J Neurosci 40, 9103–9120.

Furukawa, K., and Mattson, M.P. (1998). The transcription factor NF-kappaB mediates increases in calcium currents and decreases in NMDA- and AMPA/kainate-induced currents induced by tumor necrosis factor-alpha in hippocampal neurons. J Neurochem 70, 1876–1886.

Garcia-Oscos, F., Salgado, H., Hall, S., Thomas, F., Farmer, G.E., Bermeo, J., Galindo, L.C., Ramirez, R.D., D’Mello, S., Rose-John, S., et al. (2012). The stress-induced cytokine interleukin-6 decreases the inhibition/excitation ratio in the rat temporal cortex via trans-signaling. Biol Psychiatry 71, 574–582.

Gellner, A.K., Reis, J., Fiebich, B.L., and Fritsch, B. (2021). Electrified microglia: Impact of direct current stimulation on diverse properties of the most versatile brain cell. Brain Stimul 14, 1248–1258.

Graeber, M.B., and Streit, W.J. (2010). Microglia: biology and pathology. Acta Neuropathol 119, 89–105.

Habbas, S., Santello, M., Becker, D., Stubbe, H., Zappia, G., Liaudet, N., Klaus, F.R., Kollias, G., Fontana, A., Pryce, C.R., et al. (2015). Neuroinflammatory TNFalpha Impairs Memory via Astrocyte Signaling. Cell 163, 1730–1741.

Hagemeyer, N., Hanft, K.M., Akriditou, M.A., Unger, N., Park, E.S., Stanley, E.R., Staszewski, O., Dimou, L., and Prinz, M. (2017). Microglia contribute to normal myelinogenesis and to oligodendrocyte progenitor maintenance during adulthood. Acta Neuropathol 134, 441–458.

Heir, R., and Stellwagen, D. (2020). TNF-Mediated Homeostatic Synaptic Plasticity: From in vitro to in vivo Models. Front Cell Neurosci 14, 565841.

Hildebrandt-Einfeldt, L., Yap, K., Paul, M.H., Stoffer, C., Zahn, N., Drakew, A., Lenz, M., Vlachos, A., and Deller, T. (2021). Crossed Entorhino-Dentate Projections Form and Terminate With Correct Layer-Specificity in Organotypic Slice Cultures of the Mouse Hippocampus. Front Neuroanat 15, 637036.

Hoogendam, J.M., Ramakers, G.M., and Di Lazzaro, V. (2010). Physiology of repetitive transcranial magnetic stimulation of the human brain. Brain Stimul 3, 95–118.

Hopp, S.C. (2021). Targeting microglia L-type voltage-dependent calcium channels for the treatment of central nervous system disorders. J Neurosci Res 99, 141–162.

Jin, W.N., Shi, S.X., Li, Z., Li, M., Wood, K., Gonzales, R.J., and Liu, Q. (2017). Depletion of microglia exacerbates postischemic inflammation and brain injury. J Cereb Blood Flow Metab 37, 2224–2236.

Jurga, A.M., Paleczna, M., and Kuter, K.Z. (2020). Overview of General and Discriminating Markers of Differential Microglia Phenotypes. Front Cell Neurosci 14, 198.

Kloosterboer, E., and Funke, K. (2019). Repetitive transcranial magnetic stimulation recovers cortical map plasticity induced by sensory deprivation due to deafferentiation. J Physiol 597, 4025–4051.

Korchounov, A., and Ziemann, U. (2011). Neuromodulatory neurotransmitters influence LTP-like plasticity in human cortex: a pharmaco-TMS study. Neuropsychopharmacology 36, 1894–1902.

Laprell, L., Schulze, C., Brehme, M.L., and Oertner, T.G. (2021). The role of microglia membrane potential in chemotaxis. J Neuroinflammation 18, 21.

Lenz, M., Eichler, A., Kruse, P., Strehl, A., Rodriguez-Rozada, S., Goren, I., Yogev, N., Frank, S., Waisman, A., Deller, T., et al. (2020). Interleukin 10 Restores Lipopolysaccharide-Induced Alterations in Synaptic Plasticity Probed by Repetitive Magnetic Stimulation. Front Immunol 11, 614509.

Lenz, M., Galanis, C., Muller-Dahlhaus, F., Opitz, A., Wierenga, C.J., Szabo, G., Ziemann, U., Deller, T., Funke, K., and Vlachos, A. (2016). Repetitive magnetic stimulation induces plasticity of inhibitory synapses. Nat Commun 7, 10020.

Lenz, M., Kruse, P., Eichler, A., Straehle, J., Beck, J., Deller, T., and Vlachos, A. (2021). All-trans retinoic acid induces synaptic plasticity in human cortical neurons. Elife 10.

Lenz, M., Platschek, S., Priesemann, V., Becker, D., Willems, L.M., Ziemann, U., Deller, T., Muller-Dahlhaus, F., Jedlicka, P., and Vlachos, A. (2015). Repetitive magnetic stimulation induces plasticity of excitatory postsynapses on proximal dendrites of cultured mouse CA1 pyramidal neurons. Brain Struct Funct 220, 3323–3337.

Li, K., Wang, X., Jiang, Y., Zhang, X., Liu, Z., Yin, T., and Yang, Z. (2021). Early intervention attenuates synaptic plasticity impairment and neuroinflammation in 5xFAD mice. J Psychiatr Res 136, 204–216.

Maggio, N., and Vlachos, A. (2018). Tumor necrosis factor (TNF) modulates synaptic plasticity in a concentration-dependent manner through intracellular calcium stores. J Mol Med (Berl) 96, 1039–1047.

Masuda, T., Amann, L., Sankowski, R., Staszewski, O., Lenz, M., P, D.E., Snaidero, N., Costa Jordao, M.J., Bottcher, C., Kierdorf, K., et al. (2020). Novel Hexb-based tools for studying microglia in the CNS. Nat Immunol 21, 802–815.

Maus, L., Lee, C., Altas, B., Sertel, S.M., Weyand, K., Rizzoli, S.O., Rhee, J., Brose, N., Imig, C., and Cooper, B.H. (2020). Ultrastructural Correlates of Presynaptic Functional Heterogeneity in Hippocampal Synapses. Cell Rep 30, 3632–3643 e3638.

Muller-Dahlhaus, F., and Vlachos, A. (2013). Unraveling the cellular and molecular mechanisms of repetitive magnetic stimulation. Front Mol Neurosci 6, 50.

Muri, L., Oberhansli, S., Buri, M., Le, N.D., Grandgirard, D., Bruggmann, R., Muri, R.M., and Leib, S.L. (2020). Repetitive transcranial magnetic stimulation activates glial cells and inhibits neurogenesis after pneumococcal meningitis. PLoS One 15, e0232863.

Murugan, M., Sivakumar, V., Lu, J., Ling, E.A., and Kaur, C. (2011). Expression of N-methyl D-aspartate receptor subunits in amoeboid microglia mediates production of nitric oxide via NF-kappaB signaling pathway and oligodendrocyte cell death in hypoxic postnatal rats. Glia 59, 521–539.

Pell, G.S., Roth, Y., and Zangen, A. (2011). Modulation of cortical excitability induced by repetitive transcranial magnetic stimulation: influence of timing and geometrical parameters and underlying mechanisms. Prog Neurobiol 93, 59–98.

Pfeiffer, T., Avignone, E., and Nagerl, U.V. (2016). Induction of hippocampal long-term potentiation increases the morphological dynamics of microglial processes and prolongs their contacts with dendritic spines. Sci Rep 6, 32422.

Prinz, M., Jung, S., and Priller, J. (2019). Microglia Biology: One Century of Evolving Concepts. Cell 179, 292–311.

Prinz, M., Masuda, T., Wheeler, M.A., and Quintana, F.J. (2021). Microglia and Central Nervous System-Associated Macrophages-From Origin to Disease Modulation. Annu Rev Immunol 39, 251–277.

Riazi, K., Galic, M.A., Kentner, A.C., Reid, A.Y., Sharkey, K.A., and Pittman, Q.J. (2015). Microglia-dependent alteration of glutamatergic synaptic transmission and plasticity in the hippocampus during peripheral inflammation. J Neurosci 35, 4942–4952.

Rizzo, F.R., Musella, A., De Vito, F., Fresegna, D., Bullitta, S., Vanni, V., Guadalupi, L., Stampanoni Bassi, M., Buttari, F., Mandolesi, G., et al. (2018). Tumor Necrosis Factor and Interleukin-1beta Modulate Synaptic Plasticity during Neuroinflammation. Neural Plast 2018, 8430123.

Santello, M., Bezzi, P., and Volterra, A. (2011). TNFalpha controls glutamatergic gliotransmission in the hippocampal dentate gyrus. Neuron 69, 988–1001.

Schafer, D.P., Lehrman, E.K., Kautzman, A.G., Koyama, R., Mardinly, A.R., Yamasaki, R., Ransohoff, R.M., Greenberg, M.E., Barres, B.A., and Stevens, B. (2012). Microglia sculpt postnatal neural circuits in an activity and complement-dependent manner. Neuron 74, 691–705.

Schindelin, J., Arganda-Carreras, I., Frise, E., Kaynig, V., Longair, M., Pietzsch, T., Preibisch, S., Rueden, C., Saalfeld, S., Schmid, B., et al. (2012). Fiji: an open-source platform for biological-image analysis. Nat Methods 9, 676–682.

Sheppard, O., Coleman, M.P., and Durrant, C.S. (2019). Lipopolysaccharide-induced neuroinflammation induces presynaptic disruption through a direct action on brain tissue involving microglia-derived interleukin 1 beta. J Neuroinflammation 16, 106.

Shirinpour, S., Hananeia, N., Rosado, J., Tran, H., Galanis, C., Vlachos, A., Jedlicka, P., Queisser, G., and Opitz, A. (2021). Multi-scale modeling toolbox for single neuron and subcellular activity under Transcranial Magnetic Stimulation. Brain Stimul.

Steinmetz, C.C., and Turrigiano, G.G. (2010). Tumor necrosis factor-alpha signaling maintains the ability of cortical synapses to express synaptic scaling. J Neurosci 30, 14685–14690.

Stellwagen, D., Beattie, E.C., Seo, J.Y., and Malenka, R.C. (2005). Differential regulation of AMPA receptor and GABA receptor trafficking by tumor necrosis factor-alpha. J Neurosci 25, 3219–3228.

Stellwagen, D., and Malenka, R.C. (2006). Synaptic scaling mediated by glial TNF-alpha. Nature 440, 1054–1059.

Stevanovic, I., Mancic, B., Ilic, T., Milosavljevic, P., Lavrnja, I., Stojanovic, I., and Ninkovic, M. (2019). Theta burst stimulation influence the expression of BDNF in the spinal cord on the experimental autoimmune encephalomyelitis. Folia Neuropathol 57, 129–145.

Stowell, R.D., Sipe, G.O., Dawes, R.P., Batchelor, H.N., Lordy, K.A., Whitelaw, B.S., Stoessel, M.B., Bidlack, J.M., Brown, E., Sur, M., et al. (2019). Noradrenergic signaling in the wakeful state inhibits microglial surveillance and synaptic plasticity in the mouse visual cortex. Nat Neurosci 22, 1782–1792.

Strehl, A., Lenz, M., Itsekson-Hayosh, Z., Becker, D., Chapman, J., Deller, T., Maggio, N., and Vlachos, A. (2014). Systemic inflammation is associated with a reduction in Synaptopodin expression in the mouse hippocampus. Exp Neurol 261, 230–235.

Stroup, W.W. (2012). Generalized Linear Mixed Models: Modern Concepts, Methods and Applications. CRC Press.

Tancredi, V., D’Antuono, M., Cafe, C., Giovedi, S., Bue, M.C., D’Arcangelo, G., Onofri, F., and Benfenati, F. (2000). The inhibitory effects of interleukin-6 on synaptic plasticity in the rat hippocampus are associated with an inhibition of mitogen-activated protein kinase ERK. J Neurochem 75, 634-643.

Thimm, A., and Funke, K. (2015). Multiple blocks of intermittent and continuous theta-burst stimulation applied via transcranial magnetic stimulation differently affect sensory responses in rat barrel cortex. J Physiol 593, 967–985.

Ting, J.T., Lee, B.R., Chong, P., Soler-Llavina, G., Cobbs, C., Koch, C., Zeng, H., and Lein, E. (2018). Preparation of Acute Brain Slices Using an Optimized N-Methyl-D-glucamine Protective Recovery Method. J Vis Exp.

Vlachos, A., Muller-Dahlhaus, F., Rosskopp, J., Lenz, M., Ziemann, U., and Deller, T. (2012). Repetitive magnetic stimulation induces functional and structural plasticity of excitatory postsynapses in mouse organotypic hippocampal slice cultures. J Neurosci 32, 17514–17523.

Wang, B., Grill, W.M., and Peterchev, A.V. (2018). Coupling Magnetically Induced Electric Fields to Neurons: Longitudinal and Transverse Activation. Biophys J 115, 95–107.

Ziemann, U. (2004). TMS induced plasticity in human cortex. Rev Neurosci 15, 253–266.

Ziemann, U., Reis, J., Schwenkreis, P., Rosanova, M., Strafella, A., Badawy, R., and Muller-Dahlhaus, F. (2015). TMS and drugs revisited 2014. Clin Neurophysiol 126, 1847–1868.

