## Supplementary Information for "Microglia mediate synaptic plasticity induced by 10 Hz repetitive transcranial magnetic stimulation"

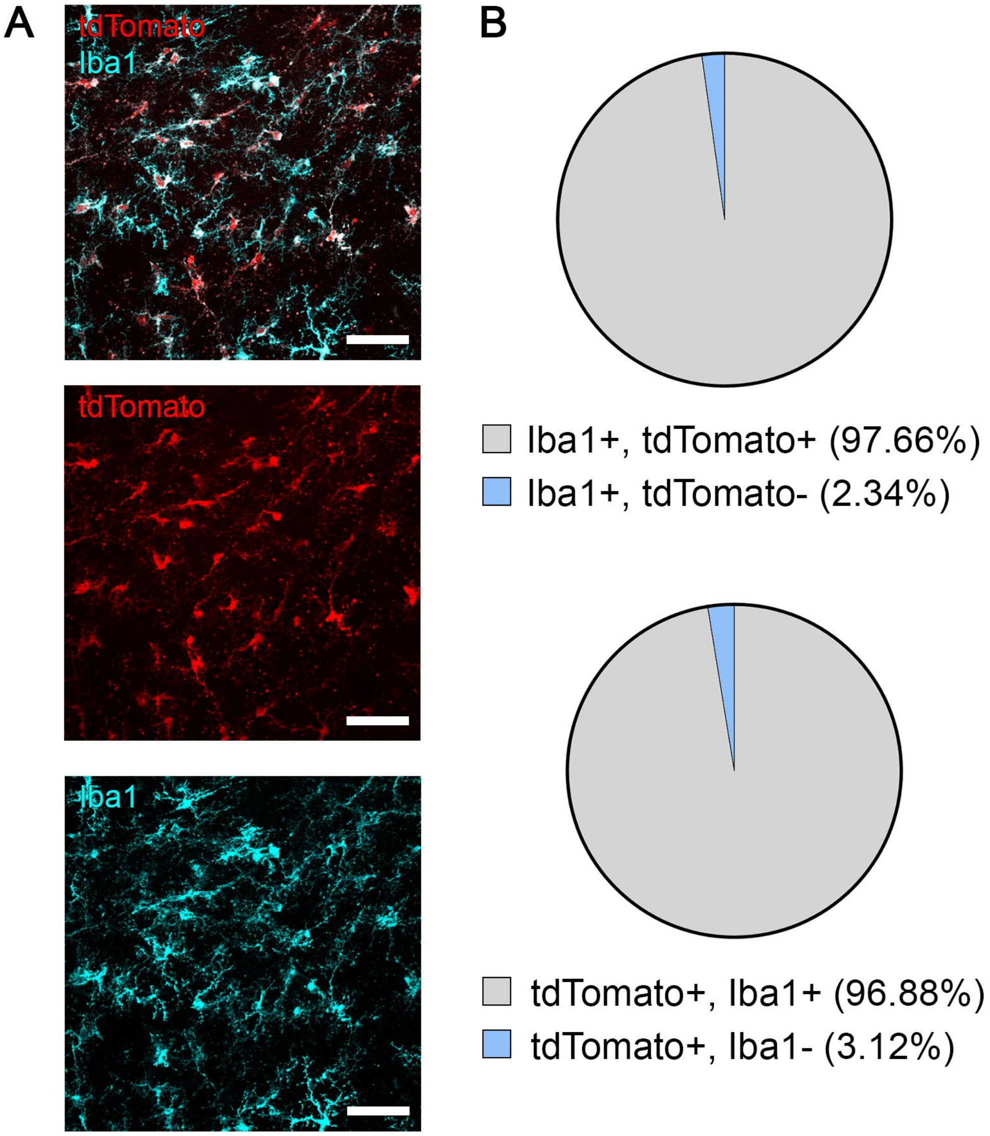


Figure S1

**Figure S1: Validation of microglia labeling in *HexB*$^{tdT/tdT}$ cultures**

(A) Representative image of Iba1 stained tissue culture prepared from homozygous *HexB*$^{tdT/tdT}$ mice. Note tdTomato expressing microglia. Scale bars, 50 µm.

(B) Group data showing an almost complete overlap of the two signals ($n_{Iba1+}$ = 273 cells in 5 cultures; $n_{tdTomato+}$ = 274 cells in 5 cultures.)

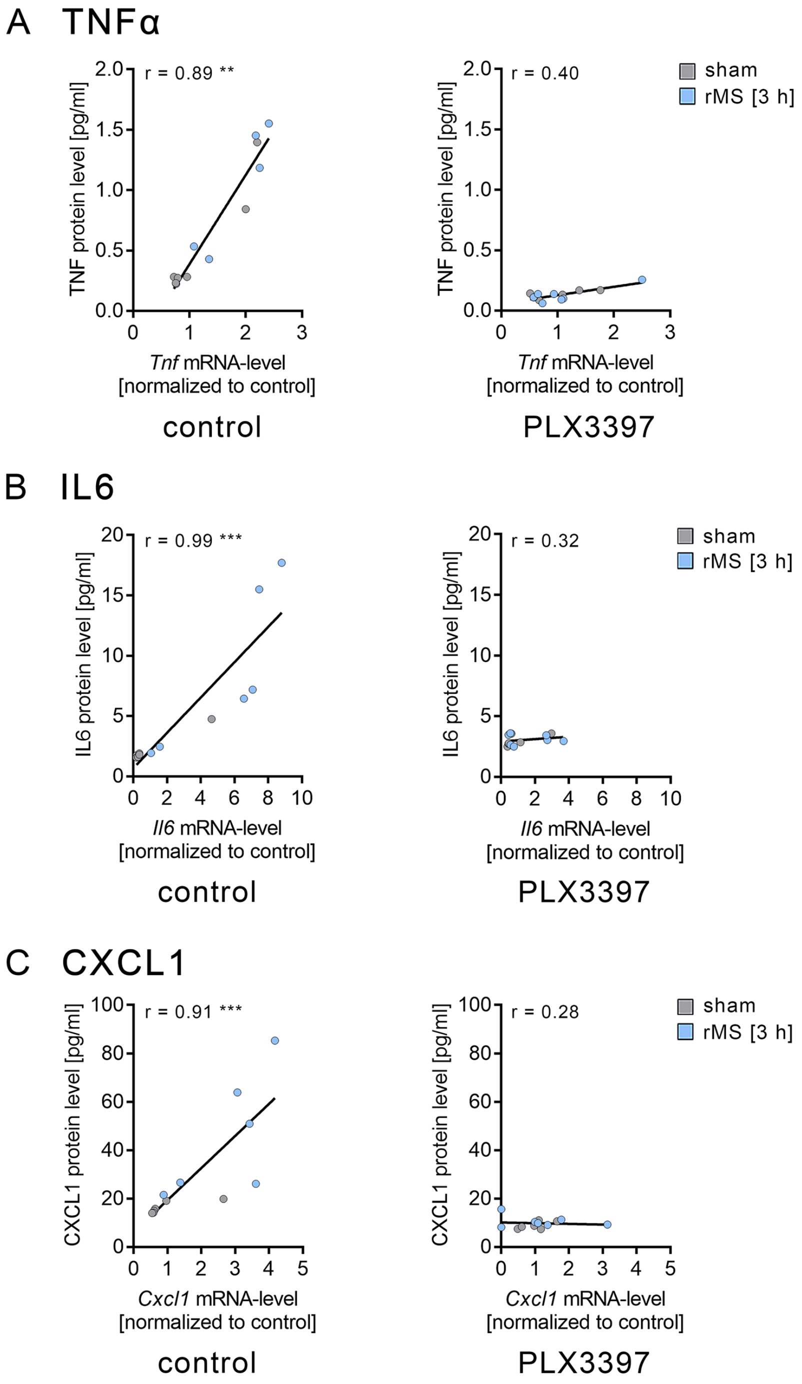


Figure S2

**Figure S2: Correlation of Cytokine mRNA and protein levels**

Correlation of mRNA levels and protein levels (separate graphs are shown in Figure 10) of TNFα (A), IL6 (B) and CXCL1 (C) in non-depleted control cultures and microglia-depleted (PLX3397 treated) cultures after sham stimulation or rMS ($n_{control}$ = 6 cultures or culturing medium samples respectively for each experimental condition, $n_{PLX3397\ sham}$ = 6 cultures or culturing medium samples, $n_{PLX3397\ rMS}$ = 7 cultures or culturing medium samples).

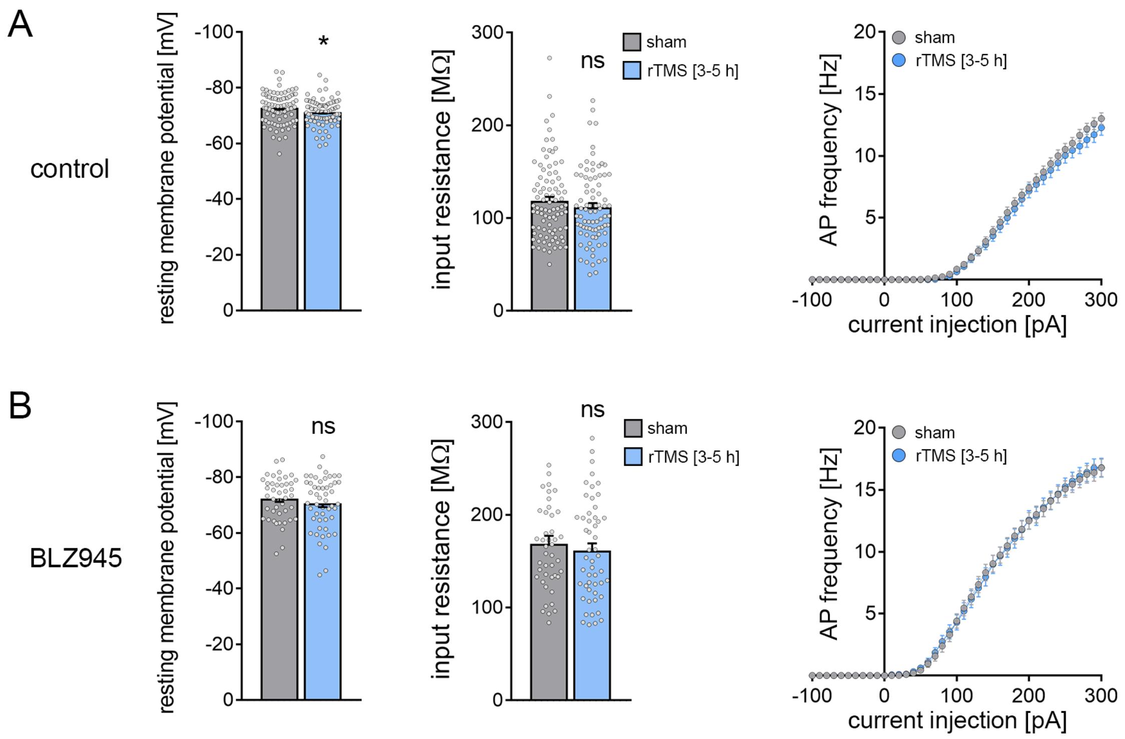


Figure S3

**Figure S3: 10 Hz-rTMS does not cause major changes in passive or active membrane properties in mPFC superficial pyramidal neurons from both depleted and non-depleted animals**

(A) Passive (resting membrane potential and input resistance) and active (action potential frequencies) membrane properties were analyzed in superficial pyramidal neurons of the mPFC from adult non-depleted mice. We found a minimal but significant increase in the resting membrane potential upon rTMS while the input resistance and the action potential frequencies remained unchanged ($n_{sham}$ = 89 cells, $n_{10Hz\ rTMS}$ = 81 cells; Mann-Whitney test and RM two-way ANOVA followed by Sidak's multiple comparisons test for action potential frequency analysis; $U_{resting\ membrane\ potential}$ = 2920).

(B) No changes in active or passive membrane properties of mPFC superficial pyramidal neurons upon rTMS were evident in microglia-depleted (BLZ945-treated) animals ($n_{sham}$ = 44 cells, $n_{10Hz\ rTMS}$ = 51 cells; Mann-Whitney test and RM two-way ANOVA followed by Sidak's multiple comparisons test for action potential frequency analysis).

Individual data points are indicated by grey dots. Values represent mean ± s.e.m (* $p < 0.05$; ns, not significant differences).

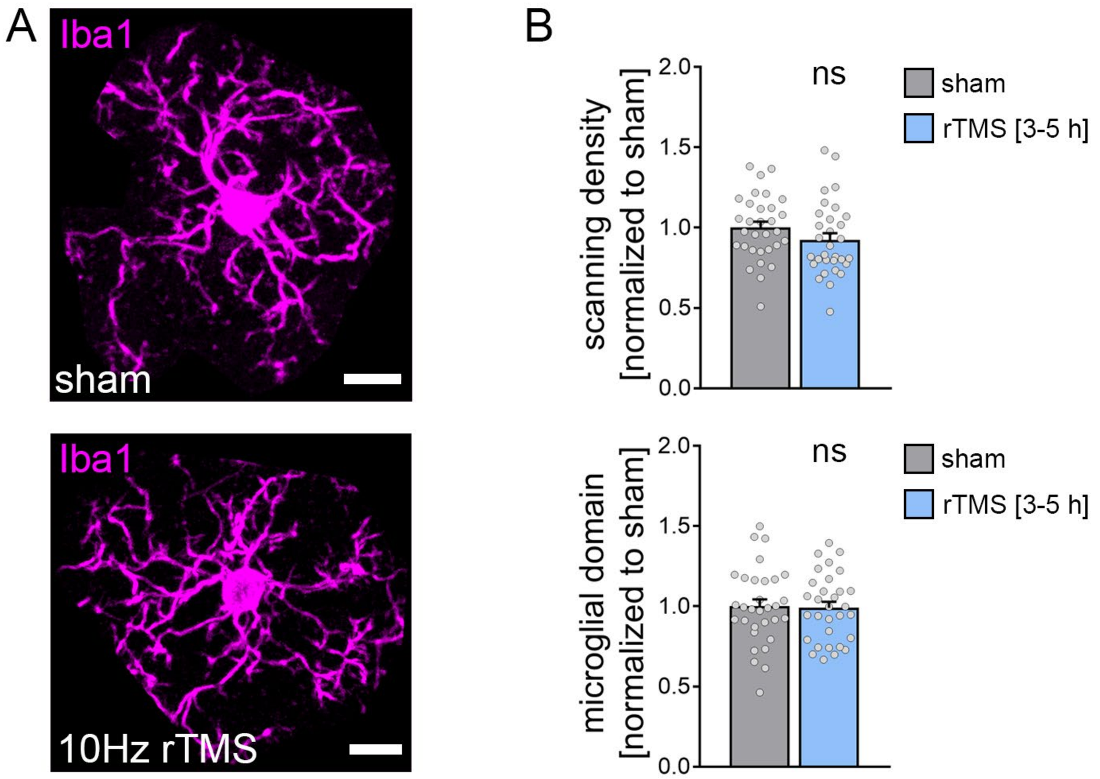


Figure S4

**Figure S4: 10 Hz-rTMS does not change morphological features of microglia in the medial prefrontal cortex of adult mice.**

(A) Representative images of Iba1-stained microglia in the medial prefrontal cortex of adult mice that have been extracted from image stacks. Microglial morphological features were compared between sham- and 10 Hz rTMS stimulated animals. Scale bars, 10 μm.

(B) No changes in both microglial scanning density and domain could be detected after stimulation ($n_{sham}$ = 31 cells from 5 animals, $n_{10Hz\ rTMS}$ = 31 cells from 5 animals; Mann-Whitney test).

Individual data points are indicated by grey dots. Values represent mean ± s.e.m (ns, not significant differences).
