## Supplementary Table 1 for "Microglia mediate synaptic plasticity induced by 10 Hz repetitive transcranial magnetic stimulation"

### Microglia mediate synaptic plasticity induced by 10 Hz repetitive transcranial magnetic stimulation

Amelie Eichler, Dimitrios Kleidonas, Zsolt Turi, Maximilian Fliegauf, Matthias Kirsch, Dietmar Pfeifer, Takahiro Masuda, Marco Prinz, Maximilian Lenz, Andreas Vlachos

| Gene Symbol | Fold Change* | P-val | FDR P-val | Description |
|---|---|---|---|---|
| Gm10600 | 3.06 | 0.0003 | 0.0444 | predicted gene 10600 [Source:MGI Symbol;Acc:MGI:3710628] |
| Gm13304 | 2.28 | 0.0001 | 0.0299 | predicted gene 13304 |
| Gm10591 | 2.22 | 0.0001 | 0.0298 | predicted gene 10591 |
| Ccl21b; Gm13304 | 2.14 | 0.0003 | 0.0484 | chemokine (C-C motif) ligand 21B (leucine); predicted gene 13304 |
| Gm13304 | 2.14 | 0.0003 | 0.0484 | predicted gene 13304 |
| Epsti1 | -2.00 | 9.13E-05 | 0.022 | epithelial stromal interaction 1 (breast) |
| Entpd1 | -2.02 | 0.0001 | 0.027 | ectonucleoside triphosphate diphosphohydrolase 1 |
| Ehd4 | -2.1 | 3.23E-05 | 0.0109 | EH-domain containing 4 |
| Plcg2 | -2.11 | 8.72E-05 | 0.0215 | phospholipase C, gamma 2 |
| Lilr4b | -2.15 | 0.0002 | 0.0381 | leukocyte immunoglobulin-like receptor, subfamily B, member 4B |
| Il10ra | -2.15 | 0.0001 | 0.0273 | interleukin 10 receptor, alpha |
| Nfam1 | -2.16 | 6.31E-05 | 0.0166 | Nfat activating molecule with ITAM motif 1 |
| Cmklr1 | -2.17 | 0.0002 | 0.0444 | chemokine-like receptor 1 |
| Cyba | -2.18 | 2.49E-05 | 0.0091 | cytochrome b-245, alpha polypeptide |
| Folr2 | -2.2 | 6.57E-05 | 0.017 | folate receptor 2 (fetal) |
| Csf3r | -2.22 | 0.0001 | 0.029 | colony stimulating factor 3 receptor (granulocyte) |
| Slfn2 | -2.24 | 7.72E-06 | 0.0049 | schlafen 2 |
| Ccr5 | -2.24 | 5.67E-05 | 0.0155 | chemokine (C-C motif) receptor 5 |
| Il1a | -2.26 | 3.04E-05 | 0.0104 | interleukin 1 alpha |
| Tmem119 | -2.28 | 0.0002 | 0.0371 | transmembrane protein 119 |
| Mylip | -2.28 | 5.62E-05 | 0.0155 | myosin regulatory light chain interacting protein |
| Cxcl16 | -2.28 | 0.0002 | 0.0319 | chemokine (C-X-C motif) ligand 16 |
| Lcp2 | -2.32 | 0.0002 | 0.0444 | lymphocyte cytosolic protein 2 |
| B430306N03Rik | -2.33 | 4.64E-05 | 0.0132 | RIKEN cDNA B430306N03 gene |
| Ikzf1 | -2.34 | 0.0001 | 0.0283 | IKAROS family zinc finger 1 |
| Olfml3 | -2.36 | 0.0001 | 0.0253 | olfactomedin-like 3 |
| Rgs10 | -2.45 | 2.10E-05 | 0.0081 | regulator of G-protein signalling 10 |
| F13a1 | -2.53 | 3.96E-05 | 0.0122 | coagulation factor XIII, A1 subunit |
| Slamf9 | -2.61 | 0.0002 | 0.0319 | SLAM family member 9 |
| Tmem86a | -2.62 | 0.0002 | 0.0381 | transmembrane protein 86A |
| Nckap1l | -2.63 | 2.13E-05 | 0.0081 | NCK associated protein 1 like |
| Mnda | -2.64 | 2.28E-05 | 0.0086 | myeloid cell nuclear differentiation antigen |

| | | | | |
|---|---|---|---|---|
| Slc2a5 | -2.65 | 0.0003 | 0.0444 | solute carrier family 2 (facilitated glucose transporter), member 5 |
| Rac2 | -2.68 | 1.49E-05 | 0.007 | RAS-related C3 botulinum substrate 2 |
| Ncf4 | -2.72 | 0.0001 | 0.0251 | neutrophil cytosolic factor 4 |
| Tyrobp | -2.77 | 6.36E-05 | 0.0166 | TYRO protein tyrosine kinase binding protein |
| Lilrb4a | -2.78 | 9.25E-05 | 0.0221 | leukocyte immunoglobulin-like receptor, subfamily B, member 4A |
| F11r | -2.78 | 7.24E-05 | 0.0185 | F11 receptor |
| Ifi30 | -2.79 | 4.51E-05 | 0.0131 | interferon gamma inducible protein 30 |
| Arhgdib | -2.79 | 3.86E-05 | 0.0121 | Rho, GDP dissociation inhibitor (GDI) beta |
| Ptafr | -2.81 | 0.0003 | 0.0484 | platelet-activating factor receptor |
| Unc93b1 | -2.83 | 1.79E-05 | 0.0076 | unc-93 homolog B1 (C. elegans) |
| Apoc2; Apoc4 | -2.87 | 0.0001 | 0.0298 | apolipoprotein C-II; apolipoprotein C-IV |
| Dock2 | -2.88 | 0.0002 | 0.0381 | dedicator of cyto-kinesis 2 |
| Cd14 | -2.9 | 9.00E-05 | 0.022 | CD14 antigen |
| Dock8 | -2.9 | 2.67E-05 | 0.0095 | dedicator of cytokinesis 8 |
| Havcr2 | -2.91 | 0.0001 | 0.0313 | hepatitis A virus cellular receptor 2 |
| Rbm47 | -2.98 | 2.69E-05 | 0.0095 | RNA binding motif protein 47 |
| Bcl2a1b; Bcl2a1a | -2.99 | 0.0002 | 0.0399 | B cell leukemia/lymphoma 2 related protein A1b; B cell leukemia/lymphoma 2 related protein A1a |
| Ncf1 | -3.04 | 1.58E-05 | 0.0072 | neutrophil cytosolic factor 1 |
| Cyth4 | -3.05 | 1.74E-06 | 0.002 | cytohesin 4 |
| Runx1 | -3.06 | 0.0001 | 0.0306 | runt related transcription factor 1 |
| Hexb | -3.1 | 1.88E-05 | 0.0078 | hexosaminidase B |
| C5ar1 | -3.13 | 1.64E-05 | 0.0073 | complement component 5a receptor 1 |
| Cd83 | -3.19 | 5.67E-05 | 0.0155 | CD83 antigen |
| Pik3cg | -3.19 | 4.54E-05 | 0.0131 | phosphoinositide-3-kinase, catalytic, gamma polypeptide |
| Vsir | -3.21 | 4.30E-05 | 0.0129 | V-set immunoregulatory receptor |
| P2ry13 | -3.22 | 0.0003 | 0.0444 | purinergic receptor P2Y, G-protein coupled 13 |
| Fcgr2b | -3.24 | 1.17E-05 | 0.0059 | Fc receptor, IgG, low affinity IIb |
| Pik3ap1 | -3.25 | 6.12E-05 | 0.0164 | phosphoinositide-3-kinase adaptor protein 1 |
| Igsf6 | -3.28 | 2.76E-06 | 0.0027 | immunoglobulin superfamily, member 6 |
| Abcc3 | -3.31 | 8.68E-05 | 0.0215 | ATP-binding cassette, sub-family C (CFTR/MRP), member 3 |
| AB124611 | -3.34 | 3.63E-06 | 0.0032 | cDNA sequence AB124611 |
| Adam8 | -3.36 | 0.0002 | 0.0381 | a disintegrin and metallopeptidase domain 8 |
| P2ry6 | -3.45 | 6.48E-06 | 0.0045 | pyrimidinergic receptor P2Y, G-protein coupled, 6 |
| Ms4a6d | -3.45 | 1.82E-06 | 0.002 | membrane-spanning 4-domains, subfamily A, member 6D |
| Ptgs1 | -3.49 | 4.92E-07 | 0.0014 | prostaglandin-endoperoxide synthase 1 |
| Tlr7 | -3.5 | 1.37E-05 | 0.0066 | toll-like receptor 7 |
| Fcgr3 | -3.53 | 3.77E-05 | 0.012 | Fc receptor, IgG, low affinity III |
| Ucp2 | -3.57 | 2.81E-05 | 0.0098 | uncoupling protein 2 (mitochondrial, proton carrier) |
| Slco2b1 | -3.64 | 2.39E-05 | 0.0088 | solute carrier organic anion transporter family, member 2b1 |

| | | | | |
|---|---|---|---|---|
| Cd180 | -3.71 | 0.0003 | 0.0448 | CD180 antigen |
| Tbxas1 | -3.75 | 1.69E-06 | 0.002 | thromboxane A synthase 1, platelet |
| Ptpn6 | -3.76 | 3.78E-05 | 0.012 | protein tyrosine phosphatase, non-receptor type 6 |
| Lcp1 | -3.76 | 1.09E-06 | 0.002 | lymphocyte cytosolic protein 1 |
| Adgre1 | -3.85 | 1.95E-06 | 0.0021 | adhesion G protein-coupled receptor E1 |
| C1qa | -3.86 | 1.57E-05 | 0.0072 | complement component 1, q subcomponent, alpha polypeptide |
| AI607873 | -3.91 | 5.76E-05 | 0.0156 | expressed sequence AI607873 |
| Csf2rb2; Mir7676-1; Mir7676-2 | -3.95 | 4.32E-06 | 0.0033 | colony stimulating factor 2 receptor, beta 2, low-affinity (granulocyte-macrophage); microRNA 7676-1; microRNA 7676-2 |
| Ncf2 | -3.95 | 0.0002 | 0.0356 | neutrophil cytosolic factor 2 |
| Ccl12 | -4.05 | 9.18E-06 | 0.0051 | chemokine (C-C motif) ligand 12 |
| P2ry12 | -4.11 | 9.76E-05 | 0.0231 | purinergic receptor P2Y, G-protein coupled 12 |
| Tlr2 | -4.11 | 7.44E-07 | 0.0018 | toll-like receptor 2 |
| Cd72 | -4.18 | 3.91E-06 | 0.0032 | CD72 antigen |
| Samsn1 | -4.19 | 3.46E-05 | 0.0115 | SAM domain, SH3 domain and nuclear localization signals, 1 |
| Tgfb1 | -4.23 | 1.69E-06 | 0.002 | transforming growth factor, beta 1 |
| Inpp5d | -4.25 | 4.87E-06 | 0.0036 | inositol polyphosphate-5-phosphatase D |
| Gpr34 | -4.41 | 1.79E-06 | 0.002 | G protein-coupled receptor 34 |
| Rasgrp3 | -4.43 | 0.0001 | 0.0313 | RAS, guanyl releasing protein 3 |
| Cd68 | -4.52 | 9.46E-06 | 0.0051 | CD68 antigen |
| Itgam | -4.57 | 1.41E-07 | 0.0006 | integrin alpha M |
| Cd53 | -4.66 | 8.94E-07 | 0.002 | CD53 antigen |
| Ly86 | -4.67 | 1.37E-05 | 0.0066 | lymphocyte antigen 86 |
| Itgb2 | -4.67 | 0.0003 | 0.0484 | integrin beta 2 |
| Plek | -4.7 | 6.60E-06 | 0.0045 | pleckstrin |
| Aif1 | -4.8 | 1.13E-05 | 0.0058 | allograft inflammatory factor 1 |
| Lpcat2 | -4.81 | 2.38E-06 | 0.0024 | lysophosphatidylcholine acyltransferase 2 |
| Cybb | -5.00 | 4.09E-08 | 0.0005 | cytochrome b-245, beta polypeptide |
| Bin2 | -5.16 | 4.00E-06 | 0.0032 | bridging integrator 2 |
| Slc15a3 | -5.17 | 1.96E-05 | 0.0078 | solute carrier family 15, member 3 |
| Slc11a1 | -5.19 | 9.00E-06 | 0.0051 | solute carrier family 11 (proton-coupled divalent metal ion transporters), member 1 |
| Laptm5 | -5.25 | 3.72E-06 | 0.0032 | lysosomal-associated protein transmembrane 5 |
| Lyz2 | -5.33 | 1.12E-07 | 0.0006 | lysozyme 2 |
| Fcer1g | -5.44 | 1.93E-05 | 0.0078 | Fc receptor, IgE, high affinity I, gamma polypeptide |
| Tnf | -5.52 | 9.67E-08 | 0.0006 | tumor necrosis factor |
| Cd37 | -5.55 | 7.98E-05 | 0.0201 | CD37 antigen |
| Fcrls | -5.59 | 1.32E-06 | 0.002 | Fc receptor-like S, scavenger receptor |
| Mpeg1 | -5.62 | 6.65E-06 | 0.0045 | macrophage expressed gene 1 |
| Selplg | -5.7 | 4.01E-05 | 0.0122 | selectin, platelet (p-selectin) ligand |
| Cd52 | -5.74 | 1.33E-08 | 0.0003 | CD52 antigen |
| Cx3cr1 | -5.75 | 3.06E-06 | 0.0028 | chemokine (C-X3-C motif) receptor 1 |
| Tlr13 | -5.76 | 1.70E-05 | 0.0074 | toll-like receptor 13 |

| | | | | |
|---|---|---|---|---|
| Pld4 | -5.91 | 1.63E-07 | 0.0006 | phospholipase D family, member 4 |
| Spi1 | -6.08 | 1.13E-06 | 0.002 | spleen focus forming virus (SFFV) proviral integration oncogene |
| Fermt3 | -6.08 | 4.39E-05 | 0.013 | fermitin family homolog 3 (Drosophila) |
| C1qb | -6.64 | 1.95E-05 | 0.0078 | complement component 1, q subcomponent, beta polypeptide |
| Trem2 | -6.93 | 1.46E-06 | 0.002 | triggering receptor expressed on myeloid cells 2 |
| Ctss | -7.08 | 3.05E-07 | 0.001 | cathepsin S |
| Gpr84 | -7.19 | 3.62E-05 | 0.0118 | G protein-coupled receptor 84 |
| C1qc | -7.53 | 8.17E-06 | 0.0049 | complement component 1, q subcomponent, C chain |
| Csf1r | -7.89 | 1.18E-06 | 0.002 | colony stimulating factor 1 receptor |
| Ccl3 | -8.34 | 7.01E-06 | 0.0046 | chemokine (C-C motif) ligand 3 |
| C3ar1 | -8.46 | 1.04E-05 | 0.0055 | complement component 3a receptor 1 |
| Spp1 | -9.64 | 8.48E-06 | 0.005 | secreted phosphoprotein 1 |
| Mmp12 | -19.27 | 8.20E-06 | 0.0049 | matrix metallopeptidase 12 |

*negative values indicate downregulation in PLX-treated tissue cultures
